# mRNA-LNP vaccination induces microenvironment remodeling and generation of tissue-resident memory T cells in glioblastoma

**DOI:** 10.64898/2026.11.11.762754

**Authors:** Shannon Snelling, Varsha Thoppey Manoharan, Joanna P. Pyczek, Xueqing Lun, Judy Wang, Jun Liu, Matthew J. Gold, Robert Nechanitzky, Kristofor Ellestad, Anushka Patel, Kirsten Olsen, Yu Wu, Yury Karpov, Haritha Menon, Yilin Tan, Rajesh Krishnan, Douglas J. Mahoney, A Sorana Morrissy, Natalia Martin-Orozco, Jennifer A. Chan

## Abstract

Glioblastoma remains highly refractory to immunotherapy, in part because of a profoundly immunosuppressive tumour microenvironment that limits effective T-cell responses. Although mRNA vaccines can induce anti-tumour immunity, how they reshape the glioblastoma immune ecosystem to support durable tumour control is not well understood. Here we use an immunocompetent EGFRvIII-driven syngeneic glioblastoma model and antigen-specific mRNA lipid nanoparticle (mRNA-LNP) vaccination to define the cellular and spatial mechanisms underlying therapeutic response. In our model, vaccination induced durable EGFRvIII-specific humoral and cellular immunity, resulting in complete tumour eradication and long-term protection against intracranial rechallenge. Integrative analyses incorporating flow cytometry, tumour interstitial fluid profiling, multiplex immunostaining, spatial transcriptomics and ligand-receptor modelling demonstrated extensive remodelling of the tumour microenvironment following vaccination. Early tumour control was associated with expansion of pro-inflammatory and antigen-presenting myeloid populations, increased cytotoxic CD8⁺ T-cell infiltration and activation, enhanced interferon-responsive signalling, and the formation of immune-supportive spatial niches enriched for antigen presentation and chemokine-mediated cellular communication. Vaccination promoted inflammatory chemokine relay networks centred on the CXCL16-CXCR6 axis, a pathway linked to tissue residency programming in the central nervous system. Functional depletion studies identified CD4⁺ and CD8⁺ T cells as essential mediators of therapeutic efficacy, while durable protection was maintained by tumour-specific brain-resident memory CD8⁺ T cells that persisted despite peripheral lymphocyte depletion. Together, these findings identify T-cell-driven microenvironmental remodelling as a critical mechanism linking mRNA vaccination to the establishment of long-term brain-resident anti-tumour immunity in glioblastoma.

## Introduction

Glioblastoma (GBM), the most common malignant primary brain tumor in adults, remains incurable, with a median overall survival of only 15 months despite multimodal treatment with surgery, radiation, and chemotherapy^1–3^. Over the past two decades, profiling of patient tumors using RNA sequencing, single-cell analyses, and spatial transcriptomics has transformed our understanding of GBM biology and has highlighted the role of the tumor microenvironment in contributing to GBM’s treatment resistance. These analyses have revealed a complex interplay between cells, where malignant cells co-opt the phenotypic plasticity of microglia, infiltrating macrophages, astrocytes, and recruited lymphoid populations to maintain a local state of immune evasion characterized by immunosuppressive myeloid populations and T-cell exclusion or dysfunction^4,5^. Interventions that overcome immunosuppression and promote stronger T-cell responses could be valuable tools in the management of this aggressive disease^6^.

Vaccination is an attractive modality for GBM because it can be tailored to generate immunity against shared or patient-specific antigens and because it can target intracellular and cell-surface antigens^7^. Clinical studies in brain tumor patients using dendritic-cell, peptide, and more recently mRNA vaccines have demonstrated that vaccine-antigen–specific humoral and cellular immunity can be generated against brain tumor neoantigens and tumor associated antigens^8,9^. However, the effects of vaccination on the GBM tumor microenvironment remain poorly characterized, owing in part to the limited availability of post-treatment tumor tissue from patients.

Here, using an immunocompetent EGFRvIII-driven syngeneic GBM model and target-specific mRNA–lipid nanoparticle vaccine, we comprehensively characterized the complex tumor microenvironmental changes underlying strong vaccine-induced brain tumor responses. Through integrated flow cytometry, immunostaining, tumor interstitial fluid analyses, and spatial transcriptomics, we found that peripherally primed T cells mediate tumor cytotoxicity within the brain. Importantly, vaccination reprogrammed the myeloid compartment from immunosuppressive states toward inflammatory and antigen-presenting phenotypes. This myeloid remodelling is associated with a TME that displays features consistent with enhanced endogenous immune surveillance and that precedes the generation of a population of tumor-specific brain tissue-resident memory (bT_rm_) T cells which sustain long-term CNS-restricted anti-tumor immunity.

## Results

### EGFRvIII mRNA–LNP vaccination elicits antigen-specific T- and B-cell responses in a syngeneic GBM model

To evaluate how tumor-specific mRNA-LNP vaccination affects the GBM tumor microenvironment, we sought an immunocompetent model with strong, uniform expression of the target gene and the ability of MHC-I to present the target. The EGFRvIII neoepitope is known to be readily presented by H-2K^k^ MHC class I molecules to CD8^+^ T cells in C3H mice^17^. We thus created C3H syngeneic GBM models by first overexpressing human EGFRvIII and knocking out CDKN2A and PTEN via brain-directed in vivo electroporation in P1 C3H pups. Cultured cells from the resulting tumors were used to establish orthotopic cell lines which we refer to as C3H-CL1-F2 (female) and C3H-CL1-M1 (male) (Supp. 1A). When orthotopically implanted into C3H/HeJ or C3HeB/FeJ mice, female syngeneic tumors arise with 100% penetrance with a median survival of 35-40 days (Supp. 1B). Syngeneic tumors recapitulate key histologic features of GBM including infiltrative growth, frequent mitoses, pseudopalisading necrosis, and vascular proliferation, and have high and uniform expression of EGFRvIII (Fig. 1A).

**Fig 1.**
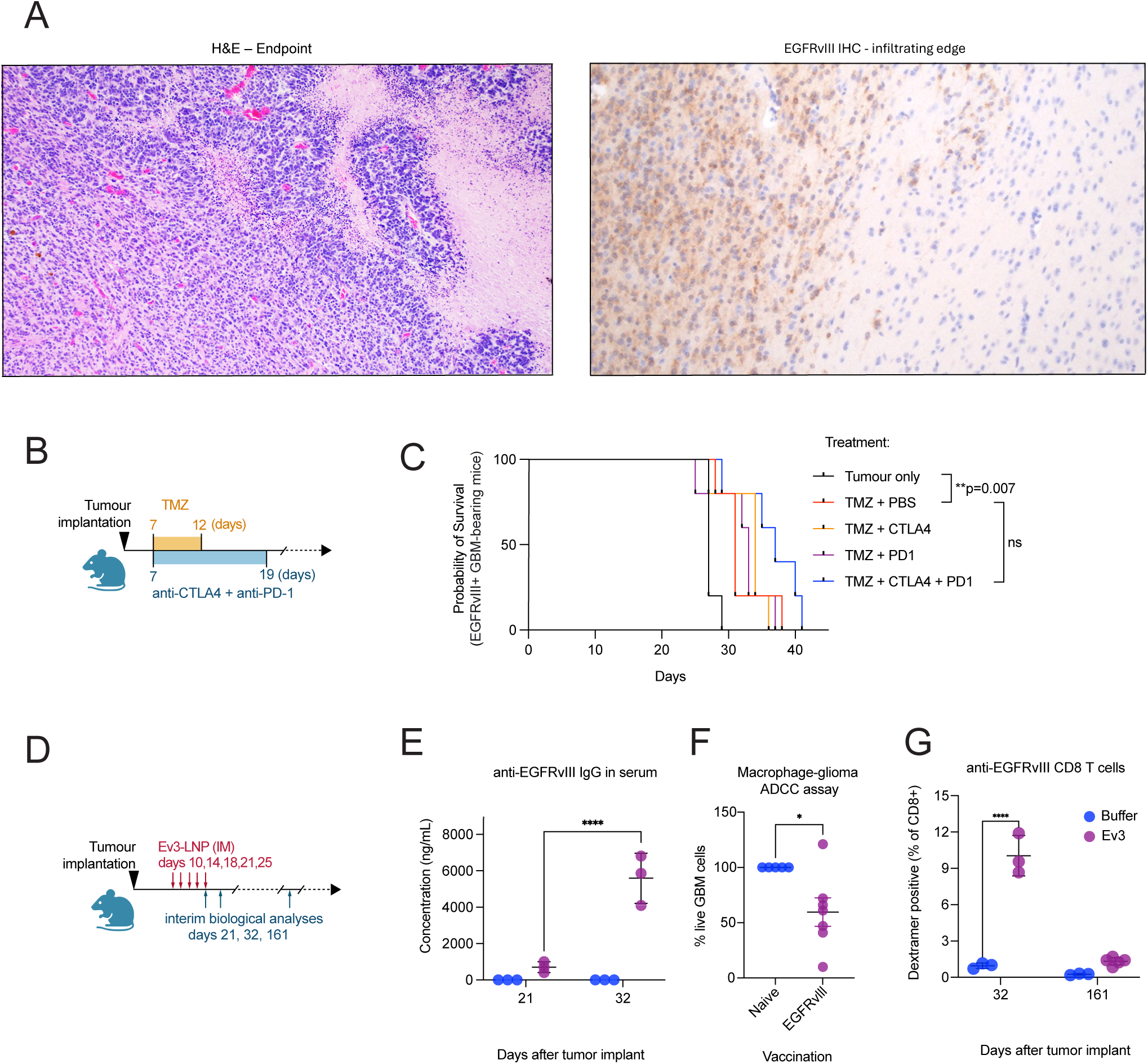
EGFRvIII mRNA-LNP vaccination elicits antigen-specific T- and B-cell responses. A) Representative H&E histology and EGFRvIII expression in the orthotopic syngeneic GBM model at endpoint. B) Experimental schematic and C) Kaplan-Meier survival analysis of the GBM-bearing mice with untreated tumor, or treatment with TMZ + PBS, TMZ + anti-PD1, TMZ + anti-CTLA-4, or TMZ + anti-PD1+ and anti-CTLA-4 therapy (n=5 mice/group; Holm-Šídák’s multiple comparisons test, ** p<0.01). D) Experimental schematic for immunogenicity testing after administration of Ev3 vaccine, luciferase control vaccine, or buffer control. Splenocytes and serum were collected at 21, 32, and 161 dpi for biological analysis (n=3-5 mice/group). E) anti-EGFRvIII antibody quantification by ELISA from serum collected at 21 and 32 dpi from Ev3 or buffer control treated (Naïve) mice. Two-way ANOVA with uncorrected Fisher’s LSD, with a single pooled variance (n=3 mice per group, ****p<0.0001). F) Antibody dependent cellular cytotoxicity/phagocytosis assay. Luciferase-expressing EGFRvIII^+^ neurospheres were co-cultured with bone marrow-derived macrophages in the presence of serum from Ev3-vaccinated animals or buffer control at 161 dpi. Target cell viability was measured by luciferase luminescence; one sample T and Wilcoxon test (n=5-7 mice/group, *p<0.05). G) anti-EGFRvIII CD8 T cells quantified by dextramer staining and flow cytometry of splenocytes at 32 and 161 dpi; one-way ANOVA with Šídák’s multiple comparisons test (n=3-5 mice/group, ****p<0.0001).

To determine whether our model recapitulates therapeutic responses observed in patients, we assessed sensitivity to current therapies. Temozolomide (TMZ) remains the standard-of-care treatment for newly diagnosed GBM, while emerging modalities such as immune checkpoint blockade (ICB) have largely failed to significantly extend survival ^2,18,19^. Consistent with clinical patterns, these models responded to but were not eradicated by TMZ (Fig. 1B-C). They also showed marginal, non-significant responses to ICB therapies (anti-PD1 and anti-CTLA4) administered alongside TMZ (Fig. 1B-C), similar to ICB resistance in the human disease.

In contrast to ICB, cancer vaccines including mRNA-LNP vaccines show promise for generating strong CD8^+^ T cell responses with anti-GBM efficacy in recent pre-clinical and clinical studies^7,20–22^. Using EGFRvIII mRNA and a proprietary LNP formulation (INTENT-2.1, WO2025010504), we generated EGFRvIII mRNA-LNPs that were uniform in size, stable, and that resulted in surface EGFRvIII expression in transfected cells (Supp. Fig. 2A). We used this EGFRvIII mRNA-LNP formulation (Ev3-LNP) to evaluate cellular and humoral response to mRNA-LNP vaccination. Tumors were initiated by orthotopic tumor cell implant and the tumor-bearing animals vaccinated at 10, 14, 18, 21 and 25 days post-tumor cell injection (dpi). Peripheral immune responses were then evaluated at 21, 32, and 161 dpi (Fig. 1D).

**Fig 2.**
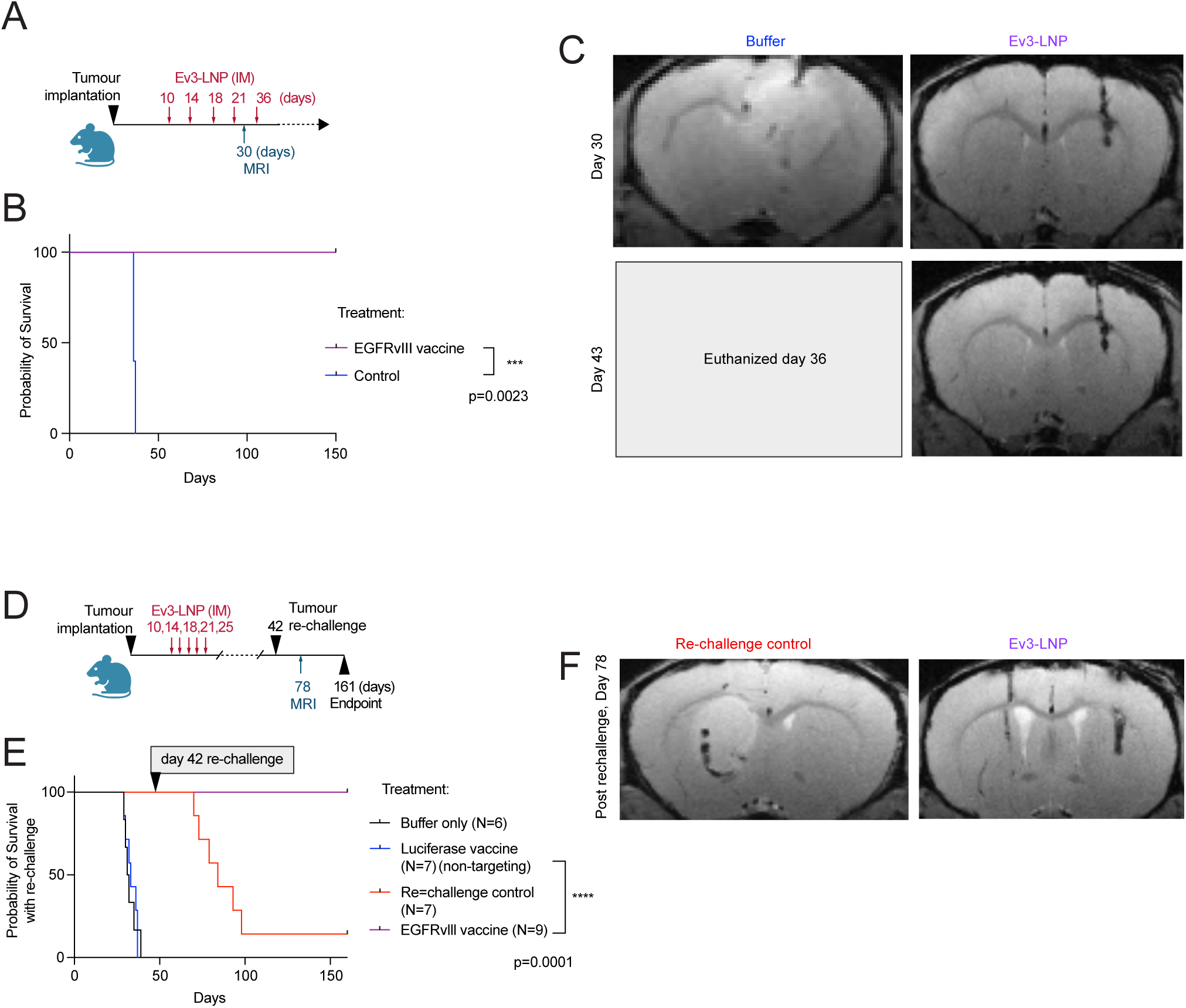
mRNA-LNP vaccination against EGFRvIII eliminates tumors and establishes durable tumor-specific immune memory. A) Experimental design and B) corresponding Kaplan-Meier survival analysis of female tumor-bearing mice treated with Ev3 vaccine or buffer control (n=5 mice/group, ***p=0.00023). C) T2 weighted MR imaging of mice at 30 and 43 dpi treated with Ev3 vaccine or buffer control. D) Experimental design and E) corresponding Kaplan-Meier survival analysis of rechallenge experiment of female tumor-bearing mice treated with Ev3 vaccine, luciferase vaccine, or buffer control. Surviving Ev3-vaccinated mice were rechallenged in the contralateral hemisphere along with a cohort of naïve age-matched mice 42 dpi (n=6-9 mice/group, ****p=0.0001). F) T2 weighted MR imaging of mice at 78 dpi previously Ev3-vaccinated or rechallenge control.

Examination of serum anti-EGFRvIII IgG titers showed that Ev3-LNP elicited humoral responses by 21 dpi (10 days after vaccine initiation), which increased further at 32 dpi (Fig. 1E). In contrast, anti-EGFRvIII antibodies were undetectable in tumour-bearing mice receiving buffer control treatment, indicating that tumour growth alone was insufficient to generate a measurable humoral response (Fig. 1E). To evaluate the potential of the humoral response to mediate anti-tumor effects, we co-cultured the serum of animals vaccinated 150 days prior with syngeneic EGFRvIII glioma cells and primary bone marrow derived macrophages for 24 hours and confirmed that vaccine-induced anti-EGFRvIII antibodies can mediate macrophage glioma-antibody dependent cellular cytotoxicity/phagocytosis (Fig. 1F).

CD8^+^ T cell response to vaccination was measured at 21 dpi by IFN-γ ELISPOT using peptide pools for human EGFR and the specific neoantigen exon 1-8 junction of human EGFRvIII, with the latter to confirm that the CD8^+^ T cells response is against the neoantigen rather than other xenogeneic differences (Supp. 2B). Using flow cytometry with an EGFRvIII H2-K^k^ dextramer, we also found evidence of target specific cellular responses induced by the vaccine. At 32 dpi (3 weeks after initiating vaccination), splenic CD8^+^ T cells recognizing EGFRvIII are significantly increased in Ev3-LNP treated animals compared to controls (mean 10.04% vs 0.96%, respectively; p<0.0001) (Fig. 1G). Furthermore, anti-EGFRvIII CD8^+^ T cells remain elevated in Ev3-LNP treated animals than controls on day 161, after last vaccination on day 25 of treatment (Fig. 1G).

Together, these findings demonstrate that Ev3-LNP induces durable humoral and cellular immunity against EGFRvIII, with antigen-specific CD8^+^ T cells persisting for more than five months after vaccination.

### mRNA–LNP vaccination against EGFRvIII eliminates tumors and establishes durable tumor-specific immune memory

To evaluate therapeutic efficacy, tumour-bearing mice received Ev3-LNP or buffer control beginning at 7 dpi and continuing through five vaccine doses (Fig. 2A). MRI performed at 30 dpi demonstrated complete radiographic tumour clearance in Ev3-LNP-treated animals, whereas substantial tumour burden was evident in control mice (Fig. 2C). Consistent with these findings, all control animals reached humane endpoint by 37 dpi, while all vaccinated mice survived without evidence of recurrence throughout the study period (150 dpi; Fig. 2B). Similar therapeutic responses were observed in an independently generated male syngeneic model, demonstrating reproducibility across sexes (Supp. Fig. 3B). Accordingly, subsequent experiments were performed in female animals.

**Fig 3.**
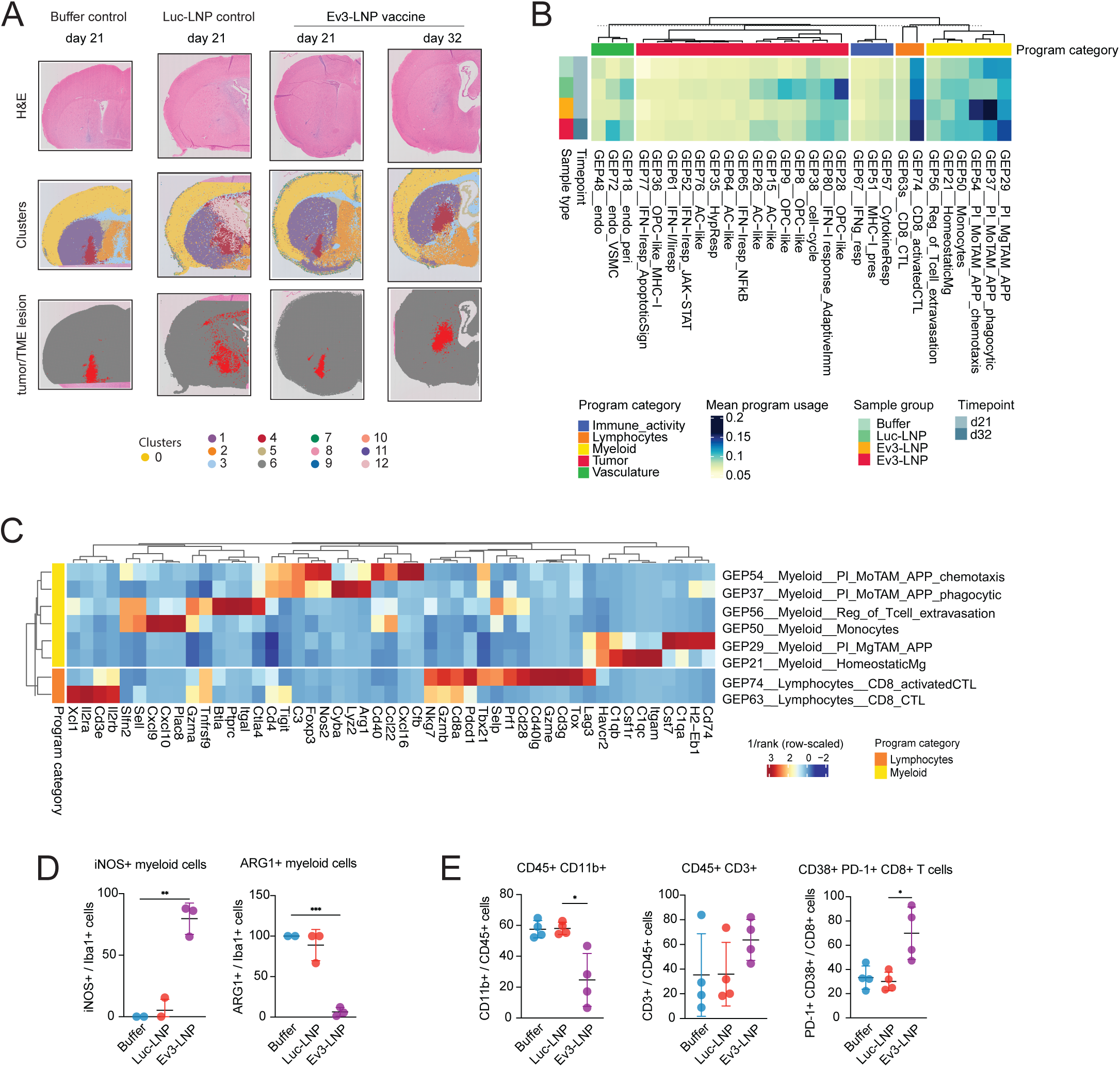
Cellular and molecular characterization of gene expression programs in VisiumHD spatial transcriptomics data. A) Spatial plots of the cohort (n = 4 samples) profiled using 10x Genomics Visium HD. Lesion (red) and non-lesion (grey) bins in the spatial plots were identified using Louvain clustering in Seurat and validated against H&E images. All quality-controlled bins (lesion and non-lesion) were subjected to consensus non-negative matrix factorization, after which lesion-enriched transcriptional programs were identified B) Hierarchically clustered heatmap showing the mean usage of lesion-enriched gene expression programs across experimental conditions. The annotation track denotes the broad cell-type or activity class assigned to each program (Myeloid, Lymphocyte, Immune activity, Vasculature or Tumor). C) Heatmap of inverse rank scores for top program-defining genes in myeloid or lymphocyte programs. D) Multiplex ARG-1, iNOS, and Iba1 immunostains of the tumor tissue of Ev3 vaccine, luciferase vaccine, or buffer control treated mice at 21 dpi (from experiment described in Fig. 2D). (n=3 mice/group, **p<0.01, ***p<0.001). E) Flow cytometry analysis of the tumor tissue of Ev3 vaccine, luciferase vaccine, or buffer control treated mice at 35 dpi with vaccination beginning at 20 dpi. (n=4 mice/group, *p<0.05).

We next asked whether vaccination established durable protective immune memory. Tumour-bearing mice were treated with Ev3-LNP, luciferase-LNP, or buffer control and surviving Ev3-LNP-treated mice were rechallenged with EGFRvIII-expressing tumour cells implanted into the contralateral hemisphere 42 days after initial tumour implantation (Fig. 2D). No additional vaccine doses were administered. Remarkably, all rechallenged mice previously treated with Ev3-LNP remained tumour-free and survived for the duration of observation (Fig. 2E). In contrast, nearly all age-matched naïve mice implanted with the same tumour cell preparation developed lethal tumours, confirming successful tumour engraftment and demonstrating that protection was specific to prior vaccination. MRI performed 36 days following rechallenge showed no detectable tumour burden in previously vaccinated mice, whereas naïve controls exhibited readily apparent intracranial tumours (Fig. 2F). Together, these findings demonstrate that mRNA-LNP vaccination not only eradicates established tumours but also generates durable immune memory capable of protecting against subsequent tumour re-exposure.

### The immune composition of the tumor microenvironment is altered by vaccination

To define how vaccination reshapes the tumour microenvironment (TME), we performed Visium HD spatial transcriptomic profiling of tumours collected from buffer-, Luc-LNP-, and Ev3-LNP-treated mice at 21 and 32 dpi (Fig. 3A). Unsupervised analysis identified 29 lesion-enriched gene expression programs (GEPs) representing distinct tumour, immune, and stromal cellular states (Fig. 3B-C, Supp. Fig. 4-6). For clarity, GEPs discussed here are referred to using both their numerical designation and an inferred biological annotation based on marker-gene expression and pathway enrichment analyses.

**Fig 4.**
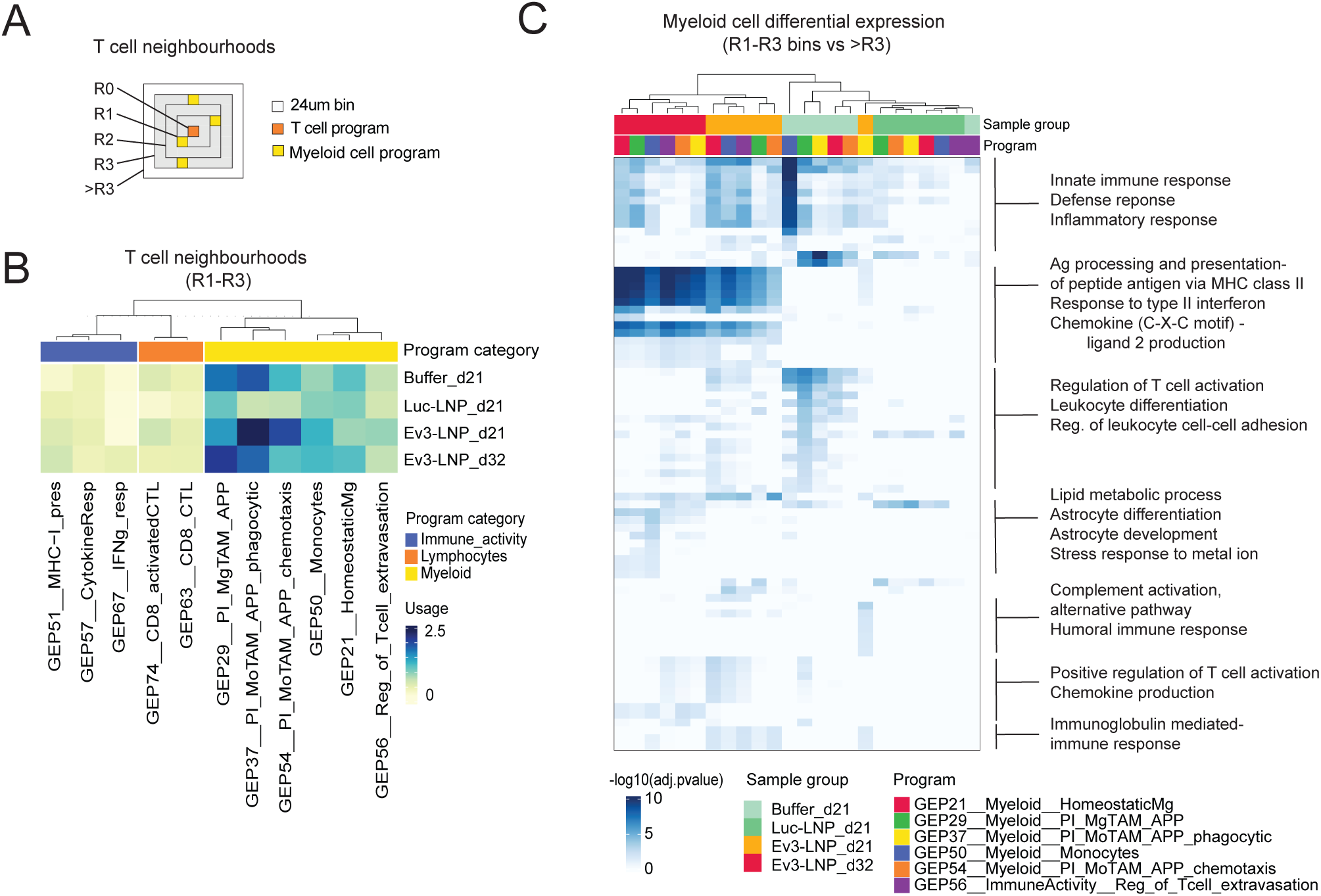
Spatial neighbourhoods of T cells. A) Schematic representation of neighbourhoods defined within radii R1-R3 of a T cell program-positive index bin (R0). B) Heatmap showing the mean usage of lesion-enriched tumor-microenvironment programs (x axis) in R1-R3 neighbours of T cell bins across experimental condition (y axis). The annotation track denotes the broad cell-type or activity class assigned to each program (Myeloid, Lymphocyte, or Immune activity). C) Pathway enrichment analysis using differentially expressed genes in myeloid program-positive bins in T cell neighbourhoods. Among the R1-R3 neighbours of the T cells, bins with > 0.05 usage of a given myeloid program were selected and tested for differential gene expression across the four experimental conditions (performed separately for each myeloid program). Significantly enriched genes per program-condition combination were subjected to GO Biological Process (GO:BP) enrichment analysis. The heatmap summarizes significantly enriched GO:BP terms (rows; adjusted p-value < 0.05), with enrichment represented as -log10(adjusted p-value).

Comparison of GEP abundance across treatment groups revealed profound remodelling of the myeloid compartment following vaccination. At 21 dpi, Ev3-LNP-treated tumours demonstrated enrichment of multiple monocyte-derived pro-inflammatory tumour-associated macrophage (PI-TAM) programs, including GEP54, GEP37, and GEP29, characterized by increased expression of genes involved in leukocyte recruitment, cytokine signalling, innate immune activation, and antigen presentation (Fig. 3B-C, Supp. Fig. 5). By 32 dpi, the antigen-presenting microglial-TAM program (GEP29) became increasingly dominant within vaccine-treated tumours (Fig. 3B, Supp. Fig. 5), suggesting a temporal transition from infiltrating inflammatory macrophages toward resident antigen-presenting myeloid populations during tumour clearance. In addition to expansion of inflammatory TAM programs, vaccination increased immune-supportive myeloid states associated with chemokine production and interferon responsiveness (GEP51, GEP57) (Supp. Fig. 5). Collectively, these findings indicate that vaccination promotes establishment of an inflammatory and immune-permissive TME capable of supporting adaptive anti-tumour immunity.

**Fig 5.**
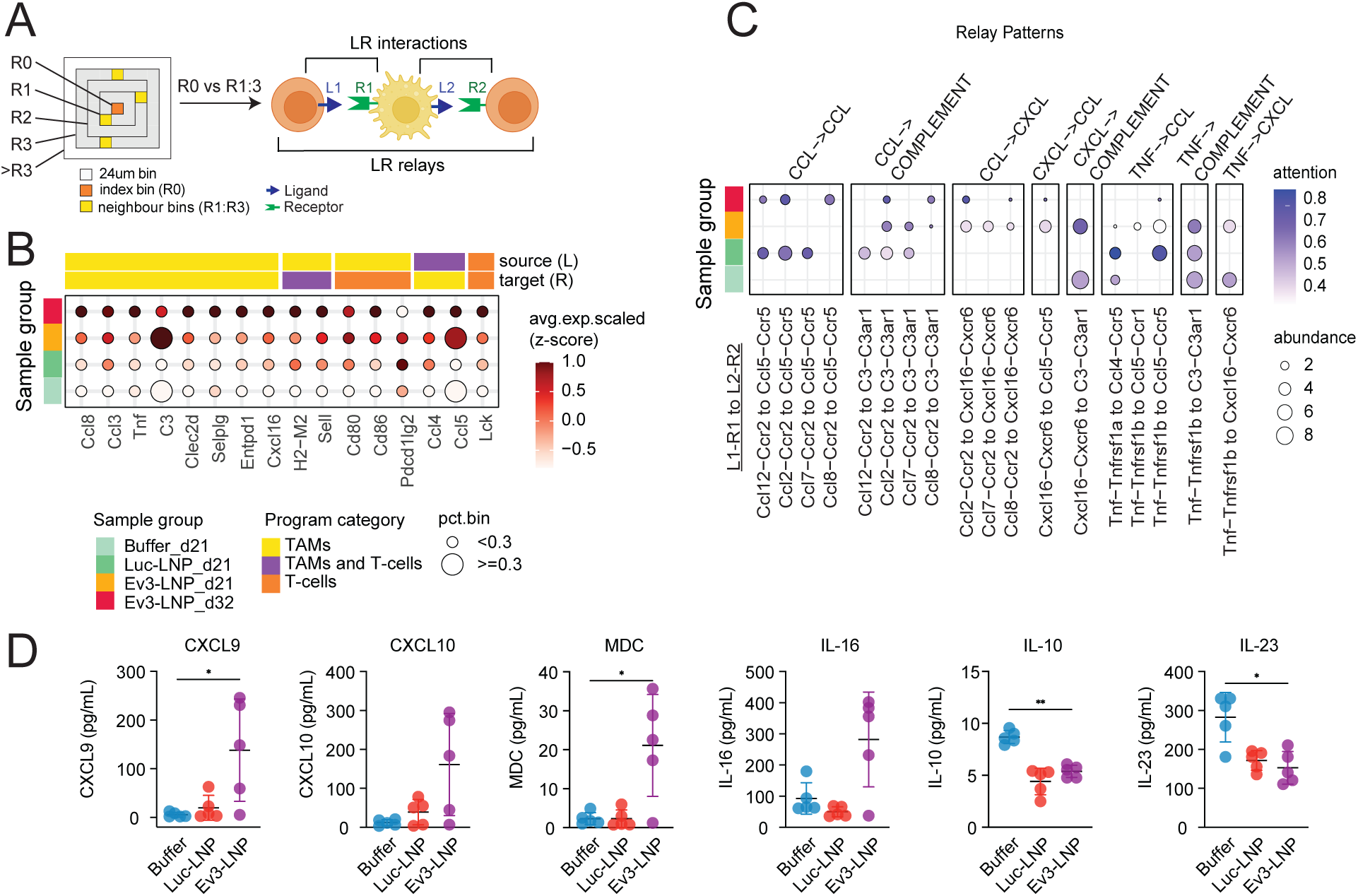
Ligand-receptor interactions across pro-inflammatory TAMs and T cells. A) Schematic outlining the workflow of ligand-receptor analysis, incorporating T cells bins at R0, and their neighbouring bins at R1-R3. B) Dotplot showing ligand expression across the four experimental conditions. Ligand-receptor pairs were filtered by retaining only those present in the top 200 genes of any lesion-enriched program, followed by selecting interactions between/within pro-inflammatory TAM and T cell programs. The source and target cell-types associated with each ligand are indicated above and below the plot, respectively. Dot size reflects the proportion of R0-R3 bins in which the ligand is expressed, and the dot color represents the expression level (z-score). C) Dotplot depicting intercellular signaling relay patterns (LR1->LR2) identified across the EGFRvIII-treated and control conditions, inferred from CellNEST. Relay patterns were defined using an edge-rank cutoff ≤200 for individual LR pairs and grouped based on the pathway of the upstream ligand–receptor pair (LR1), as annotated in the CellChat database. Dot color represents the mean attention score, and dot size indicates the abundance of the relay pattern per condition. D) Multiplexed Luminex analysis of cytokines and chemokines in the tumor microenvironment of Ev3 vaccine, luciferase vaccine, or buffer control treated mice at 35 dpi with vaccination beginning at 20 dpi (n=5 mice/group, *p<0.05 **p<0.01).

Vaccination was also associated with increased abundance of cytotoxic lymphocyte programs. While a baseline CD8 T-cell program (GEP63) was detected across all treatment groups, an activated cytotoxic T-cell program (GEP74) progressively increased following treatment and reached its highest abundance in Ev3-LNP-treated tumours (Fig. 3B-C, Supp. Fig. 5). GEP74 was characterized by expression of effector and activation-associated genes including *Gzmb*, *Ifng*, and *Lag3*, consistent with activated tumour-reactive T cells. Xenium spatial transcriptomic analysis independently confirmed increased infiltration of CD8^+^ T cells within vaccinated tumours (Supp. Fig. 7).

To validate these transcriptional findings, we performed orthogonal immunophenotypic analyses. Multiplex immunostaining demonstrated increased abundance of iNOS^+^ inflammatory myeloid cells and a reduction in Arg1^+^ immunosuppressive myeloid populations in Ev3-LNP-treated tumours relative to control groups (Fig. 3D). Similarly, flow cytometry confirmed remodelling of the tumour immune compartment, with a relative reduction in myeloid populations and an increase in T cells, including an increase in CD38^+^PD-1^+^CD8^+^ T cells consistent with enhanced CD8 T-cell activation following vaccination (Fig. 3E). Together, these analyses demonstrate that Ev3-LNP vaccination induces coordinated activation of inflammatory myeloid and cytotoxic T-cell programs within the GBM microenvironment.

### T cell–secreted interferon-γ drives remodeling of spatial immune neighbourhoods and signaling networks in the brain

To determine how these immune populations were spatially organized within the tumour microenvironment, we analyzed cellular neighbourhoods surrounding cytotoxic T-cell programs. Vaccination markedly altered the composition of these regions, with T-cell-rich niches becoming increasingly enriched for pro-inflammatory antigen-presenting TAM programs and interferon-responsive states. Notably, the dominant TAM population associated with T-cell neighbourhoods shifted from monocyte-derived inflammatory macrophages at 21 dpi toward microglia-derived antigen-presenting macrophages at 32 dpi, consistent with dynamic evolution of local immune niches during tumour eradication (Fig. 4A-B, Supp. Fig. 8B-C). Pathway analyses further revealed enrichment of antigen processing, antigen presentation, and type II interferon (IFNγ) signaling within T-cell-associated neighbourhoods of vaccinated tumours (Fig. 4C, Supp Fig 9), indicating that vaccination promotes localized immune ecosystems optimized for T-cell effector activity.

Given their close spatial proximity, we next investigated intercellular communication between proximal CTLs and PI-TAMs by inferring ligand-receptor interactions (Fig. 5A; Supp. Fig. 10A). We found that T cell-TAM ligand-receptor interactions in Ev3-LNP treated tumors were characterized by T-cell co-stimulatory signaling through the CD80/CD86–CD28 axis, as well as regulatory signaling mediated by the PD-1–PD-L1 pathway. In addition, we observed ligand–receptor pairs involved in T-cell recruitment such as Ccl3/8 – Ccr2/5, immune synapse formation and T-cell recruitment, including Cxcl16–Ccr6, Cxcl16-Cxcr6 interactions and multiple MHC class II–associated antigen presentation signals (Fig. 5B; Supp. Fig. 10B). Beyond direct LR inference, we considered longer *LR relays* whereby a receptor (R1) engaged by an initial ligand (L1) leads to increased expression of a second ligand (L2) on the receiving cell, that interacts with a second receptor (R2) (see conceptual schematic in Fig 5A). This analytic strategy inferred a chemokine relay specific to Ev3-LNP treated lesions, in which Ccl2/7/8-Ccr2 signaling in PI-TAMs induced Cxcl16 expression, which in turn engaged Cxcr6 on PI-TAMs and CTLs (Fig. 5C) – suggesting the emergence of inflammatory chemokine relay networks capable of reinforcing T-cell recruitment and residency. Since the Cxcl16-Cxcr6 axis has been shown to orchestrate brain CD8^+^ T cell tissue residency and spatial co-localization with microglia in the CNS^23^, this observation suggests that CTLs within Ev3-LNP-treated lesions may be undergoing niche-directed tissue residency programming.

We quantified cytokines and chemokines in tumor interstitial fluid at the protein level and observed increased concentrations of inflammatory mediators associated with lymphocyte recruitment—including CXCL9, CXCL10, and MDC in Ev3-LNP-treated mice compared with controls (Fig. 5D). Additionally, EGFRvIII vaccine-treated tumors showed reduced levels of the immunosuppressive cytokine IL-10 compared with controls (Fig. 5D).

### Brain-resident memory CD8^+^ T cells mediate long-term protection following vaccination

Local tissue-specific cues and inflammation determine the final differentiation of effector lymphocytes into different long-lived lineages. We aimed to understand which protective adaptive immune responses were mediating long-term immunity following local vaccine-induced changes in the tumor microenvironment of Ev3-LNP vaccinated mice.

To determine which lymphocyte immune subsets were required to confer vaccine protection, we depleted CD8^+^ T cells, CD4^+^ T cells, or B cells from EGFRvIII^+^ tumor bearing mice prior to and during vaccination (Fig. 6A). Mice depleted of CD4^+^ or CD8^+^ T cells at the time of vaccination succumbed to tumor burden (Fig. 6A), whereas mice depleted of B cells survived, suggesting that T cell immunity is required in our model. We hypothesized that EGFRvIII specific CD8^+^ T cell responses in the brain were required for survival, and that CD4^+^ T cell depletion in the periphery limited the generation of this response because of the requirement for CD4 “help” in cytotoxic T cell responses. To measure this population in the brain, we conducted flow cytometry on tumors during the depletion following three vaccine doses in these mice on 21 dpi (Fig. 6A). As expected, the frequency of CD8^+^ T cells in the tumor CD45 compartment decreased in animals peripherally depleted of CD4^+^ or CD8^+^ T cells compared to isotype control (Fig. 6B). Within the tumor’s residual CD8^+^ T cell compartment, the absolute numbers of EGFRvIII-specific CD8+ T cells were also markedly reduced, though not entirely absent, indicating that a sufficient quantity of tumour-specific CD8^+^ T cells is required for tumour clearance.

**Fig 6.**
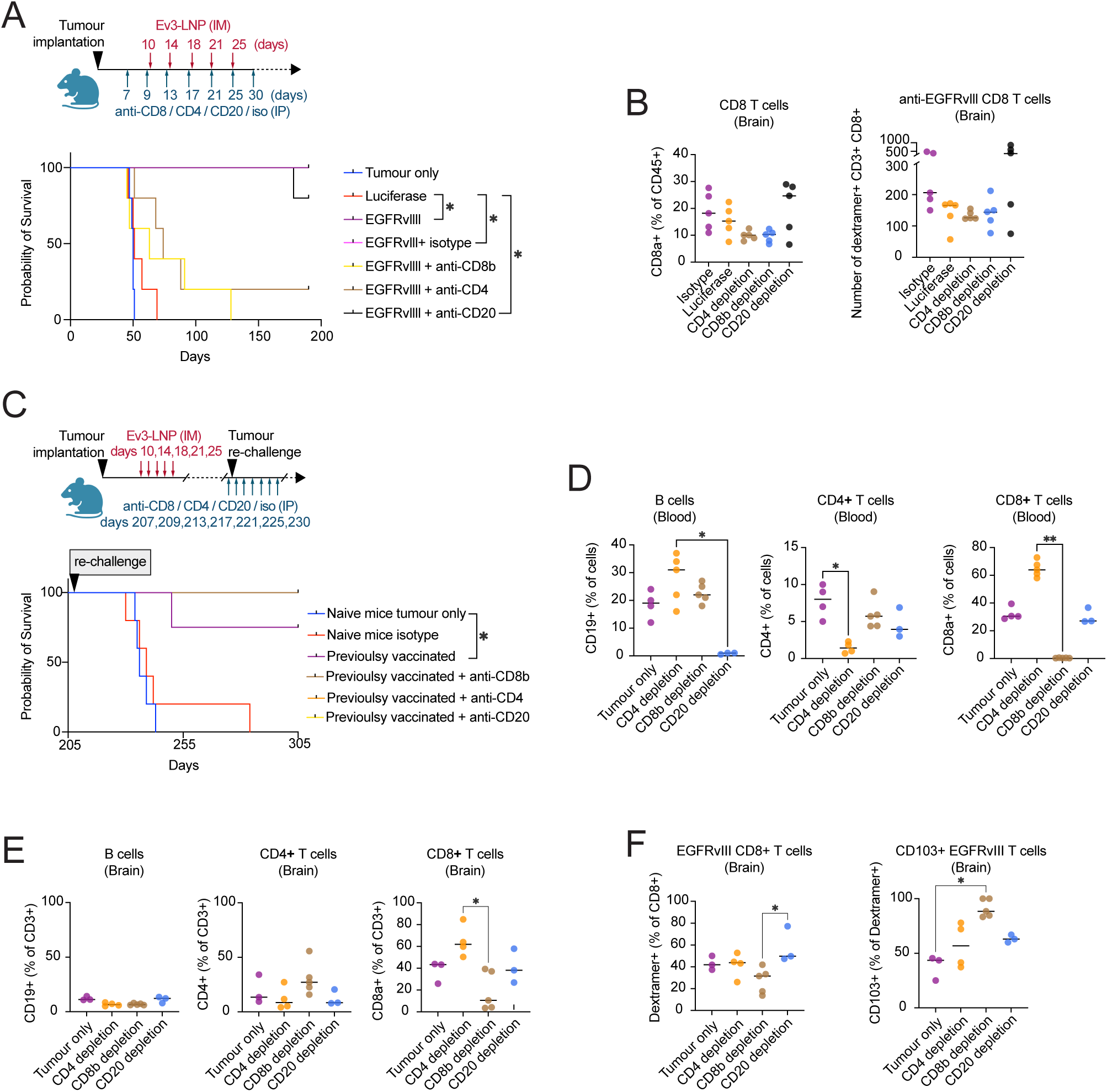
Brain-resident memory CD8 T cells mediate long-term protection after vaccination. A) Experimental design of survival experiment with CD4^+^, CD8^+^, or CD20^+^ lymphocyte depletion or isotype control prior to and during vaccination and corresponding Kaplan-Meier survival analysis (n=5 mice/group, *p<0.05). B) Flow cytometry analysis of the tumor tissue at 21 dpi from Ev3 vaccine-, luciferase vaccine-, or buffer control-treated mice with lymphocyte depletion depicted in panel A (n=4 mice/group, *p<0.05). C) Experimental design of survival experiment with lymphocyte depletion and corresponding Kaplan-Meier survival analysis after late tumor re-challenge, at over 200 dpi (n=5 mice/group, *p<0.05). D) Flow cytometry analysis of the peripheral blood of Ev3 vaccine treated mice (depleted of lymphocytes with antibodies as in Fig. C) following depletion (n=5 mice/group, *p<0.05). E) Flow cytometry analysis of the tumor tissue of Ev3-vaccinated mice (depleted of lymphocytes with antibodies as in Fig. C) 24 hours following recall challenge. (n=3-5 mice/group, *p<0.05). F) anti-EGFRvIII CD8^+^ T cell dextramer staining and flow cytometry analysis of the tumor tissue of Ev3 vaccinated mice (depleted of lymphocytes with antibodies as in Fig. C) 24 hours following recall challenge with 0.5e6 cells into the striatum. (n=3-5 mice/group, *p<0.05).

To understand which immune cell subset persists in the brain in the long-term after vaccination and mediates protection from recurrence, we vaccinated EGFRvIII^+^ tumor bearing mice and then rested them for 200 days prior to depleting CD8^+^ T cells, CD4^+^ T cells, or B cells from the animals and then challenging them again with EGFRvIII^+^ tumor (Fig. 6C). Validation of the peripheral depletion confirmed almost complete depletion of CD8^+^ T cells, CD4^+^ T cells, or B cells in the blood (Fig. 6D). Peripheral depletion of CD8^+^ T cells, CD4^+^ T cells, or B cells did not prevent the survival of rechallenge animals compared to isotype control (Fig. 6C). We suspected that vaccine induced bT_rm_ CD8^+^ T cells were mediating protection against rechallenge and that these cells reside in CNS niches which were not accessed by peripherally administered depletion antibodies. To confirm this hypothesis, we challenged the previously vaccinated animals depleted of CD8^+^ T cells, CD4^+^ T cells, or B cells a third time with a bolus of 0.5x10^6^ EGFRvIII^+^ tumor cells in the original injection hemisphere (day 305) and then conducted spectral flow on the lymphocytes in the injection hemisphere of the brain at 24 hours to evaluate the rapid recall response of CD8^+^ bT_rm_ cells. Though the frequencies of CD8^+^ T cells in the brain were reduced compared to non-CD8 depleted groups, CD8^+^ T cells and EGFRvIII-specific CD8^+^ T cells were detected (Fig. 6D-E). Moreover, most of these CD8 depletion-resistant EGFRvIII-specific CD8^+^ T cells expressed CD103, an integrin associated with resident T cell populations, suggesting that vaccination-induced bT_rm_ protect animals from rechallenge in our model (Fig. 6F).

## Discussion

Our integrated spatial transcriptomic and protein-level analyses support a model in which early activation of anti-EGFRvIII T cells remodels the tumor microenvironment, driving polarization of the myeloid compartment toward an inflammatory, anti-tumor phenotype. This shift reduces immunosuppressive signalling and enhances subsequent T-cell recruitment through chemokine and cytokine release, as well as via direct immune synapse formation and MHC-mediated antigen presentation. Together, these processes amplify T-cell effector function over time and ultimately support the establishment of long-term bT_rm_ anti-tumor T cells.

In a physiologically relevant syngeneic EGFRvIII GBM model, we show that mRNA–LNP vaccination induces potent and durable antigen-specific CD8^+^ T-cell and B-cell responses capable of mediating complete tumor clearance and long-term protection against rechallenge. Vaccination efficacy depends on T-cell immunity, while durable protection is maintained by tumour-specific brain-resident memory CD8+ T cells that persist despite extensive depletion of circulating lymphocyte populations. Spatial transcriptomic, protein-level, and immune-phenotyping analyses demonstrate that activation of vaccination-generated T cells remodels the GBM tumor microenvironment by driving inflammatory myeloid polarization, reducing immunosuppression, enhancing T-cell recruitment, and restructuring immune neighbourhoods toward anti-tumor signaling networks. Together, these data define a mechanistic framework in which T-cell–driven remodeling of the tumor microenvironment precedes the emergence of protective bT_rm_ cells and durable anti-tumor immunity following mRNA-LNP vaccination.

BT_rm_ cells are a distinct lineage of T cells which develops following pathology and rapidly responds to antigen re-encounter in the brain without re-circulating outside of the central nervous system (CNS)^24^. Among the T cell phenotypes which could be generated for GBM immunotherapy, bT_rm_ cells are particularly well suited to sustain anti-tumour immunity over the long term. CD8^+^ bT_rm_ have been demonstrated to retain high TCR affinity in the brain, making them very sensitive to antigen^25^. In addition, bT_rm_ cells rapidly remodel the brain TME in syngeneic mouse models of pathology by activating and recruiting peripheral immune cells and endogenous microglia^26^. Considering the prevailing opinion in the field that success of immunotherapy in GBM is predicated on ability to overcome immunosuppression, our finding that mRNA-LNP vaccination is able to promote the development of bT_rm_ cells in the immune environment is notable.

By showing how vaccination restructures these immune communities and myeloid activation, our work also provides a mechanistic rationale for combination therapies designed to overcome suppressive myeloid programs and reinforce productive T-cell responses. These insights offer a framework for pairing vaccines with other immunotherapies that depend on permissive immune landscapes, such as bispecific antibodies, TCR-engineered T cells, or CAR-T therapies. Vaccination is an especially compelling therapeutic platform for GBM because it is modular and capable of targeting diverse antigen classes, including neoantigens or overexpressed self-antigens^27^. In contrast to engineered cell therapies such as CAR-T cells which often fail to persist in the CNS^28^, vaccination may provide a more physiologic and sustainable avenue for generating brain-resident T cells. This distinction and potential complementarity has significant implications for therapeutic application.

Limitations to this study and model must be acknowledged. First, although our model preserves an intact tumor microenvironment, it does not incorporate human HLA restriction, which limits direct translation to human T-cell epitope recognition. Second, EGFRvIII is not an ideal clinical monotherapy target due to antigen heterogeneity^9^ and although we employed it as a proof-of-principle antigen, future studies should incorporate multi-antigen vaccines to address tumor escape using heterogenous models. Third, achieving complete immune depletion within the CNS and in niches outside the blood remains technically challenging, which may underestimate the contribution of different lymphocyte populations. Lastly, syngeneic models typically exhibit rapid tumor kinetics, which may selectively favour fast-acting T-cell–dependent therapies and underrepresent the kinetics of B-cell or antibody-mediated processes.

Finally, the relevance of our findings depends on preclinical models that recapitulate human therapeutic responses. Early mouse studies of immune checkpoint blockade failed to predict human inefficacy in GBM^29^, highlighting the need for models with appropriate immunosuppressive microenvironments. The ability of our model to recapitulate both responsiveness to standard therapy and resistance to immune checkpoint blockade strengthens confidence that the observed mechanisms of vaccine-induced TME remodelling may be relevant to human GBM.

## Methods

### Ethics approvals

All animal experiments were performed at the University of Calgary and approved by the University of Calgary Animal Care Committee (protocols AC21-0228 and AC25-0002).

### Cell lines

PiggyBac transposon/transposase and CRISPR-Cas9 were used to overexpress human EGFRvIII and knock out endogenous CDKN2A and PTEN in neural progenitors in P1 postnatal pups via *in vivo* electroporation into lateral periventricular cells. Resulting tumors were used to generate the C3H-CL1-F2 (female) and C3H-CL1-M1 (male) cell lines. Syngeneic GBM cells were cultured as neurospheres in NeuroCult Basal Medium (Mouse & Rat) with NeuroCult Proliferation Supplement (StemCell Technologies 05702) with additional 2ug/ml heparin (StemCell 07980), 20ng/ml FGF (StemCell 78003.2), and 20ng/ml EGF (StemCell 78006.2). Cells were maintained in 5% CO2 at 37 °C. Lines were routinely authenticated by driver antigen expression.

### Animal models and tissue processing

Female C3H/HeJ (Jackson Labs, strain #:000659) or C3HeB/FeJ mice (Jackson Labs, strain #:000658) age 6-8 weeks were implanted orthotopically with 10^5^ syngeneic tumor cells in 2uL of PBS by stereotactic injection into the striatum (2.0–3.0 mm right of bregma, 1.0 mm anterior to coronal suture). Mice were followed for clinical symptoms and were euthanized at either pre-determined experimental timepoints or at humane endpoint. Blood was processed for serum. Brains and spleens were harvested for histology or fresh tissue dissociation. Tissue for histology was fixed for 24 hours in 10% neutral buffered formalin and processed for paraffin embedding. For viable cells, tumor tissue was dissociated using the Tumor Dissociation Kit, mouse (Miltenyi - 130-096-730) on a gentleMACS Octo Dissociator (Miltenyi) per manufacturer’s protocol. Dissociated cells were frozen in CELLBANKER 2 (Amsbio 11914) for later analyses.

### In vivo treatments

mRNA-containing LNP vaccines were prepared using the proprietary INTENT LNP platform (Providence Therapeutics, WO2025010504). Briefly, nucleotide modified mRNAs were transcribed *in vitro* using T7 RNA polymerase, purified and encapsulated with INTENT-2.1 LNP. mRNAs included human EGFRvIII (Ev3-LNP) and full-length luciferase mRNA (luc-LNP) as a non-targeting control, or buffer only control. All mRNA LNPs passed the release tests of Providence Therapeutics, including particle size, polydispersity index, encapsulation efficiency, mRNA integrity, mRNA identity, pH, osmolality, and bacterial endotoxin. Vaccines or controls were delivered intramuscularly into the thigh approximately every 3 days for 5 doses of 10ug each, beginning at 7 or 10 days post-tumor implant (dpi). For temozolomide (Sigma-Aldrich T2577), drug was administered daily for 5 days beginning at 7 dpi, at a dose of 50 mg/kg by oral gavage. For checkpoint inhibitors, anti-PD-1 clone RMP1-14 (Bio X Cell BE0146) and anti-CTLA-4 clone 9D9 (Bio X Cell BE0164) were delivered as 200ug every 3 days for 5 doses by intraperitoneal injection, starting at 7 days post-implant.

### anti-EGFRvIII antibody quantification

Recombinant human EGFRvIII protein (Sino Biological) was coated to MaxiSorp ELISA plate (ThermoFisher) via overnight incubation. The plate was then washed with TBST (ThermoFisher) and blocked with 3% BSA for 1 hour at room temperature. Mouse serum samples diluted in 0.1% BSA were then added to the plate. Serially diluted mouse anti-human EGFRvIII mAb (Sigma) was also added to the plate as standard control. After 2.5 hours of incubation at room temperature, the plate was washed with TBST and incubated with HRP-labelled goat anti-mouse IgG (H+L) (BioRad). The plate was then washed with TBST and incubated with TMB substrate (ThermoFisher) for 11 min at room temperature. Stop solution (ThermoFisher) was added to the plate and OD450 values were measured in an ELISA reader (Promega). OD450 values of the mouse sera samples were converted to mass concentration using the standard curve generated from the OD 450 values of the serially diluted mouse anti-human EGFRvIII mAb.

### Antibody-dependent cellular cytotoxicity (ADCC) assay

Target C3H-CL1-F2 cells with luciferase overexpression were dissociated with Accumax (Innovative Cell Technologies, Inc. - AM105), mechanically triturated, and resuspended in warmed NeuroCult medium (see *Cell lines* section above) before being counted and plated at defined densities (10,000–100,000 cells) in 96-well plates. Effector macrophages derived from bone marrow progenitors and polarized to the macrophage lineage for 9 days in M-CSF (50ng/mL) in RPM1+ 10% FBS were harvested and resuspended at 1×10⁶ cells/mL prior to addition to target cells at a 1:1effector-to-target ratio. Serum from previously vaccinated animals at day 161 post-tumor cell implant or buffer was added, and plates were incubated for 24 hours at 37 °C. Cell viability was assessed by adding luciferin substrate, incubating for 5 minutes, and measuring luminescence on a plate reader (Molecular Devices, SpectraMax iD3). Luminescence values were recorded and exported for downstream analysis.

### ELISPOT

Splenocytes were plated into ELISPOT plates (MABTECH) and stimulated for 24 hours by culturing in media alone or in media containing pooled EGFRvIII overlapping peptides (1μg/mL per peptide), or PMA/Ionomycin (positive control) (250,000 cells/well). Biotinylated anti-IFN-γ antibody and streptavidin-HRP (diluted in PBS with 0.5% FBS) were used for detection, with TMB as the substrate for visualization following manufacturer’s instructions. Overnight-dried plates were imaged on an S6 Entry M2 ImmunoSpot analyzer (Cellular Technology Limited) and quantified with ImmunoSpot 5.1.34 software.

### Dextramer Staining

Two million splenocytes or tumor infiltrating immune cells from tumor tissues were thawed and stained with LIVE/DEAD fixable dye (1:1000), followed by Fc block (1:100) in FACS buffer. Cells were resuspended in EGFRvIII dextramer stain solution consisting of 10uL MHC dextramer H-2 Kk/EEKKGNYV – PE (Immunodex) in 50uL of FACS buffer for 20 minutes at room temperature. For experiments including multiple stains, the *spectral flow cytometry* method was preformed next (see below). After washing, fixing and permeabilizing, cells were resuspended in 300uL of FACS buffer. Data was acquired on an Aurora 5L (CyTEK).

### MR imaging

MR imaging of isoflurane-anaesthetized animals was performed with a 9.4T Bruker Avance system with 3.5 cm diameter quadrature coil (Bruker) using a multi-slice, T2-weighted sequence and a single T1 slice through the middle of the tumor. The field of view was 2-3 cm^2^, slice thickness was 0.75 mm, and the data matrix was 256x256 or 128x128. T2 maps were acquired using a multi-slice multi-echo spin-echo sequence (e.g. TR = 2500 ms, TE = 8 ms, 25 echos, 4 averages). T1 parameters were TR/TE = 4.2/1.8 ms and 12 inversions with 12 different T1 delays between 234 and 9226 ms for the inversion pulses.

### Spectral flow cytometry

2 million thawed cells were stained with a LIVE/DEAD aqua stain (Thermofisher L34965) followed by Fc blocking (Biolegend 156603). Cells were stained with an extracellular antibody mix for 20 minutes, followed by fixation/permeabilization (eBioscience 00-5521-00) and intracellular antibody staining as previously described^4^. Antibodies used are detailed in Supplementary Table 1. Flow cytometry was performed on an Aurora 5L (CyTEK).

### Immunohistochemistry

5 µm thick FFPE sections on Superfrost Plus slides (VWR) were immunostained per routine protocol. Antigen-retrieval was performed in pH 9 Antigen Retrieval Solution (Akoya Biosciences) in the microwave (2 minutes at full power, then 20 minutes at 20% power). Peroxidase block (Dako) followed by blocking buffer (Dako) were used prior to primary antibody incubation. Primary antibody diluted in Antibody Diluent (Cell Signaling) was applied for 1 hour at room temperature. Secondary polymer HRP (Rabbit, Dako) was applied for 30 minutes at room temperature. Staining visualized with DAB reagent (Dako) per manufacturer’s protocol, with hematoxylin as a counterstain. Slides were coverslipped with Epredia mounting media. Primary antibody details are provided in Supplemental methods.

### Multiplexed immunostaining

OPAL multiplex immunofluorescence staining (Akoya Biosciences) was performed per manufacturer’s protocol. For each OPAL staining cycle, slides were heated in pH 9 Antigen Retrieval Solution in the microwave, as above. Primary antibody was diluted in Antibody Diluent (Cell Signaling) and incubated on slides for 1 hour at room temperature. Secondary polymer HRP was incubated on the slides for 10 minutes at room temperature. OPAL reagent (1:200 in diluent) was added to slides for 10 minutes. After washing, the steps from antigen retrieval though OPAL reagent application were repeated for each staining cycle. Spectral DAPI in TBS-T was incubated on the slides for 5 minutes. Slides were washed with TBS-T and coverslipped with FluorSave mounting media.

### Visium HD library preparation, sequencing and preprocessing

*Tissue preparation and library construction.* FFPE brain tissue from four mice (Buffer_d21, Luc-LNP_d21, Ev3-LNP_d21 and Ev3-LNP_d32) were sectioned coronally at 5 um onto plain glass slides. Sections were deparaffinized, H&E stained and imaged at 40x magnification on a VS110 slide scanner (Olympus), following recommendations in the Visium HD FFPE Tissue Preparation Guide (CG000684). Sections were then hybridized with the Visium Mouse Transcriptome Probe Set v2.0 supplemented with custom EGFRvIII probes. Using the CytAssist instrument (software v2.1.0.15), probe products were transferred to the 6.5 x 6.5 mm capture areas of Visium HD slides. Probe extension and library construction followed the Visium HD Spatial Gene Expression Reagent Kits User Guide (CG000685). Libraries were sequenced on an Illumina NextSeq 2000 with paired-end reads (Read 1, 43 cycles; i7 and i5, 10 cycles each; Read 2, 50 cycles), and FASTQ files were generated onboard with DRAGEN BCL Convert (v4.2.7).*Custom probe design for EGFRvIII detection*: To enable the detection of EGFRvIII transcripts in tissue sections profiled using 10x Genomics Visium HD spatial transcriptomics, nine custom probes (50 base pairs each; 25 bp LHS and 25 bp RHS probes) targeting the EGFRvIII coding sequence were designed in accordance with 10x Genomics guidelines for custom probe design. Probe sequences were generated using custom code available on GitHub and complete sequences are provided **in** Supplementary Table 2.

### Xenium data analysis

10x Genomics Xenium profiling was performed on coronal brain sections from six samples representing buffer control, Luc-LNP, and Ev3-LNP conditions at day 21 and day 32 dpi, using a predesigned 247-gene panel supplemented with 100 custom genes. Xenium Onboard Analysis output was imported into R using Seurat (v5.1.0), with cell and transcript coordinates linked to their corresponding field-of-view (FOV) images for each of the six tissue sections. Cells with fewer than 10 detected transcripts were excluded (nCount_Xenium > 10). Per-sample objects were merged with unique cell-barcode prefixes and normalized using SCTransform on the Xenium assay across the full gene panel. Principal component analysis was performed on the SCT-normalized data using all genes in the panel, and the first 50 principal components were integrated across samples using Harmony to correct for sample-to-sample batch effects. A shared nearest-neighbor graph was constructed using the first 30 Harmony-integrated dimensions, followed by Louvain clustering at a resolution of 0.5. Marker genes per cluster were identified using the Wilcoxon rank-sum test on SCT-normalized data using FindAllMarkers(). Genes with a Bonferroni-adjusted p < 0.05 and average log2 fold-change > 1 were considered significant markers. T-cell, monocyte-derived TAM (Mo-TAM), and microglia-derived TAM (Mg-TAM) clusters were annotated based on differentially expressed genes and used to assess cell-type proportions across treatment conditions and time points.

### Alignment and Quantification

Raw sequencing data were processed using Space Ranger (version 3.1.2) for read alignment and gene count matrix generation. A custom reference genome was constructed by integrating the mouse reference genome (mm10/GRCm38) with EGFRvIII construct sequences using *spaceranger mkref* with default parameters. For samples buffer_d21 and Ev3-LNP_d21, which contained secondary tissue sections on the slide, manual image alignment was performed using Loupe Browser (version 9.0.0) to exclude non-primary tissue regions prior to quantification. FASTQ files were then aligned to the custom reference using *spaceranger count* with a bin size of 24 μm and default parameters otherwise. The resulting gene expression matrices were imported into R and processed using Seurat (version 5.1.0^10^). Initial quality control retained bins expressing at least one gene and genes detected in at least one bin. The filtered gene expression matrices from all four samples (Buffer_d21, Luc-LNP_d21, Ev3-LNP_d21 and Ev3-LNP_d32) were merged using the *merge()* function in Seurat to create a unified dataset for downstream analysis.

### Visium HD data analysis

#### Identification of gene expression programs

Transcripts captured within each Visium HD bin (24 μm) may originate from multiple cells with distinct identities or activity states, necessitating deconvolution to resolve the underlying cellular contributions and gene signatures. To this end, consensus non-negative matrix factorization (cNMF), an unsupervised deconvolution method, was applied to decompose the merged dataset into distinct gene expression programs (GEPs) without relying on pre-defined cell-type signatures^11^. To select features representing informative biological variation, over-dispersed (OD) genes were identified using the *restrictCorpus()* function from the R package STDeconvolve (version 1.3.2^12^). Genes were filtered to retain those detected in more than 1% but fewer than 100% of bins, followed by selection of OD genes with a significance threshold of α = 0.05. To ensure comprehensive coverage of biological variation across samples, OD genes were identified independently for each condition as well as for the aggregated dataset, and the union of all OD gene sets was taken, resulting in a total of 1825 genes. Bins that did not express at least one OD gene were removed from further analysis. The aggregated filtered raw count matrix and the list of OD genes were provided as input to cNMF and factorization was performed across a range of ranks (k = 10 - 100) with Frobenius norm as the beta-divergence loss function for 200 iterations per rank (n_iter = 200). Consensus programs were generated per rank using a density threshold of 0.1 during the consensus clustering step to ensure robust and reproducible program identification.

### Rank selection and lesion identification

To determine an optimal rank (k) for downstream analyses, each factorized cNMF solution was evaluated based on two criteria: (1) the number of informative programs, defined as programs with usage >5% in at least 5% of bins, and (2) the number of lesion-enriched programs. To identify lesion regions, unsupervised clustering was performed using Seurat on the merged dataset. The top 10 principal components computed from the normalized gene expression matrix were utilized for KNN graph-based clustering with Louvain algorithm at a resolution of 0.1, which yielded a distinct cluster corresponding to the lesion. This assignment was validated by concordance with EGFRvIII probe expression patterns and H&E histology. Lesion-enriched programs were then identified as those with significantly higher usage within lesion bins compared to non-lesion bins (two-sided Wilcoxon rank-sum test; log2 fold-change > 1; Benjamini-Hochberg adjusted p-value < 0.05). Based on these criteria, rank k=80 was selected, which maximized both the number of informative programs and lesion-enriched programs (Supplementary Table 3).

### Usage normalization and thresholding

Program usage values from the k = 80 solution were normalized such that the sum of usage values across all programs equaled 1 for each bin. A bin was considered to express a given program if its normalized usage exceeded 0.05. This threshold was consistently applied for program assignment and for calculating mean program usage within lesion bins across conditions.

### Annotation of Gene Expression Programs

Lesion-enriched gene expression programs were annotated using two methods as previously described^13^: (1) marker gene scoring to assess similarity to known cell-type and cell-state gene signatures curated from multiple studies of the mouse brain, GBM, and the GBM TME, and (2) Gene Ontology Biological Process (GO:BP) enrichment analysis^13^. The marker gene scores yielded unambiguous assignments for cell-type associated programs, while pathway enrichments validated the classification of programs into broad cellular categories (immune, vascular, brain and tumor) and identified distinct biological activities associated with individual programs.

### Spatial coincidence networks

Directed weighted graphs were used to construct (1) a global network (GN), quantifying spatial co-localization (SC) between lesion-enriched programs across the cohort representing their global spatial organization, and (2) differential networks (DN) to identify condition-specific changes in spatial organization, as previously described^13^. Both networks were visualized using igraph (v2.0.3), with nodes representing individual programs and directed edges pointing from query (i) to target (j), with edge width scaling proportionally to coincidence magnitude (SC*_i_*_→*j*_ or ΔSC *_i_*_→*j,*_ for GN and DN, respectively). In the GN, the "graphopt" layout was used, with edges drawn when SC*_i_*_→*j*_ > 20%, node size reflecting total program-positive bins across the aggregated data, and node color indicating assigned cell-type or activity class. In the DN, node positions were fixed to the GN layout to enable direct comparison, with edges retained if |ΔSC*_i_*_→*j*_| ≥ 0.20 and Fisher’s exact p < 0.05. Edge color encodes direction of change (red: increased; blue: decreased), node size reflects the number of samples with ≥50 bins at usage >0.1, and node color encodes the symmetric log2 fold-change in program-positive bin proportions between conditions. Together, GN summarizes cohort-wide co-localization patterns among lesion-enriched programs, while the DN highlights condition-specific alterations, revealing changes in cellular spatial communication.

### Spatial Neighborhood Analysis and Differential Expression

#### T Cell Neighborhoods

To investigate the cellular composition surrounding T cells, T cell-enriched bins were first identified as those expressing GEP63 or GEP74 (T cell programs; normalized usage >0.05) within lesion bins. For each T cell-enriched bin, the spatial neighborhood was defined as all bins within a 3-bin radius (approximately 96 μm). The mean usage of each lesion-enriched program within these neighborhood bins was then calculated separately for each experimental condition and results were visualized as a heatmap.

### Characterization of myeloid programs in T cell neighborhoods

To understand the functional characteristics of myeloid programs in proximity to T cells in the lesion, pathway enrichment analysis was performed on genes differentially expressed in myeloid-enriched bins within T cell neighbourhoods. For each myeloid program within each experimental condition, bins in T cell neighbourhoods expressing the program (normalized usage > 0.05) were identified and compared to all other T cell neighbourhood bins using the FindAllMarkers() function in Seurat with the Wilcoxon rank-sum test^10^. Differentially expressed genes were defined as those with Bonferroni adjusted p-value < 0.05 and log2 fold-change >1. Marker genes identified for each condition-myeloid program combination were then subjected to GO:BP enrichment analysis using g:Profiler (version 0.2.3^14^) with the g:SCS multiple testing correction algorithm. Enrichment was assessed using the hypergeometric test, with only GO terms containing between 10 and 2000 genes included. Results were visualized as a heatmap of -log10(adjusted p-values) to identify biological processes associated with myeloid populations in the T cell microenvironment across conditions.

### Ligand-Receptor interaction analysis

To identify ligand-receptor (LR) interactions mediating cellular communication between myeloid populations and T cells, spatially aware LR analysis was performed using CellChat (version 2.1.2) with the mouse ligand-receptor database^15^. Analyses were restricted to T cell-enriched bins within lesion regions (GEP63 or GEP74 usage >0.05) and their local spatial neighborhoods (all bins within a 3-bin radius). These bins were annotated as “T cells” and “neighbours”, respectively, and used as the two interacting groups for the analysis. CellChat was run independently for each condition with the following parameters: communication probability estimation using truncated mean (type = "truncatedMean", trim = 0.01), distance-weighted interactions enabled (distance.use = TRUE, interaction.range = 100 μm, scale.distance = 0.05), and contact-dependent signaling parameters (contact.dependent = TRUE, contact.range = 48 μm, contact.knn.k = 20). Statistically significant ligand-receptor pairs (p < 0.05) were identified based on permutation testing within CellChat. To focus on program-level communication, LR pairs were filtered to retain only those where both the ligand and receptor genes ranked within the top 200 genes (ranking based on cNMF gene spectra z-scores) of any lesion-enriched program. Signaling directionality was then inferred by assigning each ligand and receptor gene to the program in which it achieved its highest rank, thereby identifying source (ligand-expressing) and target (receptor-expressing) programs. Interactions were subsequently filtered to include only those occurring between or within myeloid and T cell programs. For these program-level LR pairs, the mean inverse rank (1/rank) of the ligand gene in the source program and the receptor gene in the target program was computed and visualized as a heatmap, with higher values indicating stronger gene-program associations. In addition, normalized expression levels of ligand genes across the four experimental conditions were visualized using dot plots.

To resolve relay networks of TAM-T cell communication, CellNEST was applied to VisiumHD gene expression data restricted to tumor periphery bins with LR pairs sourced from the CellChat mouse database^16^. Preprocessing was performed using --filter_min_cell=1, --threshold_gene_exp=98, and kNN-based spatial neighborhoods (k=50), with default juxtacrine and neighborhood distance thresholds of 139.66 and 558.64 pixels, respectively, encompassing communication within three bin hops. The model was trained once per sample using a fixed random seed (seed=1) and default architecture parameters (60,000 epochs, hidden dimension=512). Post-processing retained the top 20% of interactions by attention score, from which relay networks were identified. Networks were subsequently restricted to those containing at least one CellChat-shortlisted LR pair, and interactions with edge_rank < 200 were retained to prioritize high-confidence communication events. Mean attention scores were then computed per LR pair and averaged across both constituent LR pairs in a relay network to yield a single representative score. To ensure directional program-level continuity, only networks in which the target program of LR1 matched the source program of LR2 were retained. Final relay networks were subset to those identified in Ev3-LNP-treated conditions, grouped by LR1 signaling pathway as defined by the CellChat database and visualized as dot plots, with dot size and color encoding network abundance and mean attention score, respectively.

### Statistical Analyses

Statistics for non-spatial transcriptomic experiments were performed using Prism software 10.1.1 (GraphPad). Spatial transcriptomic data (Visium HD and Xenium) were analyzed in R v4.3.1 and Python v3.12.2 using the packages described above. Comparisons between two groups were performed using the two-sided Wilcoxon rank-sum test, and p-values were adjusted for multiple comparisons using the Benjamini-Hochberg method for usage comparisons and the Bonferroni method for differential expression analysis. Adjusted p-values < 0.05 were considered statistically significant.

## Data and code availability

Raw and processed VisiumHD spatial transcriptomics data are deposited in the Gene Expression Omnibus (GEO accession GSE352858). Processed Xenium data and data to reproduce figures from spatial transcriptomic analyses are available at Zenodo (10.5281/zenodo.22257399), and the corresponding code is available on GitHub (https://github.com/MorrissyLab/Ev3_mRNA-LNP_vaccine__VisiumHD).

## Contributions

Conceptualization: JAC, ASM, NMO; Methodology: SS, VTM, JPP, KE, DJM, JL, MG, ASM, JAC, NMO; Investigation: SS, JPP, XL, AP, JW, RN, HM, YT; Resources: DJM, KO, YW, YK, RK; Formal analysis: VTM, SS; Data Curation: ASM; Writing – original draft: SS, VTM; Writing – review & editing: JAC, ASM, NMO; Supervision: JAC, DJM, ASM, NMO, JL

## Supplementary Figures

**Supplementary Figure 1.**
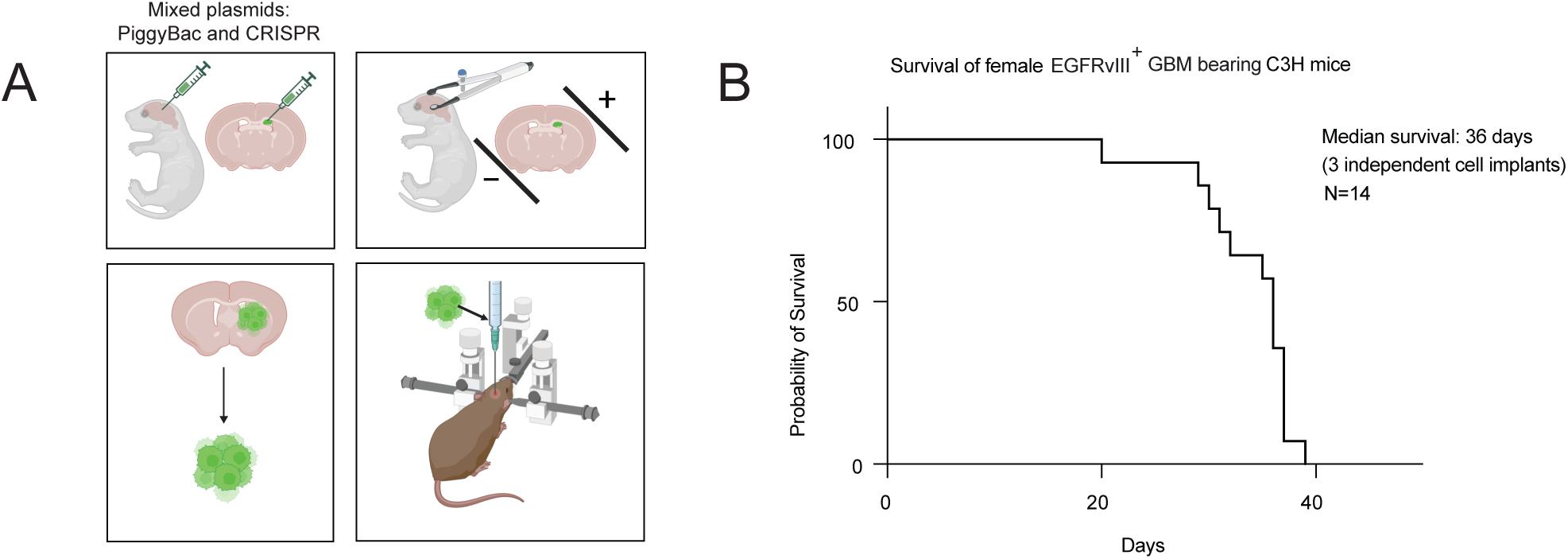
Generation and baseline characterization of the syngeneic EGFRvIII+ GBM model. A) Schematic of model generation. Human EGFRvIII was overexpressed and endogenous Cdkn2a and Pten knocked out in lateral periventricular neural progenitors of P1 C3H pups by in vivo electroporation of PiggyBac transposon/transposase and CRISPR-Cas9 plasmids. Resulting tumors were cultured as neurospheres to establish the C3H-CL1-F2 (female) and C3H-CL1-M1 (male) lines, which were implanted orthotopically into syngeneic recipients. B) Kaplan-Meier survival of female C3H mice bearing orthotopic EGFRvIII+ tumors (median survival 36 days; n = 14 mice pooled across three independent cell implants).

**Supplementary Figure 2.**
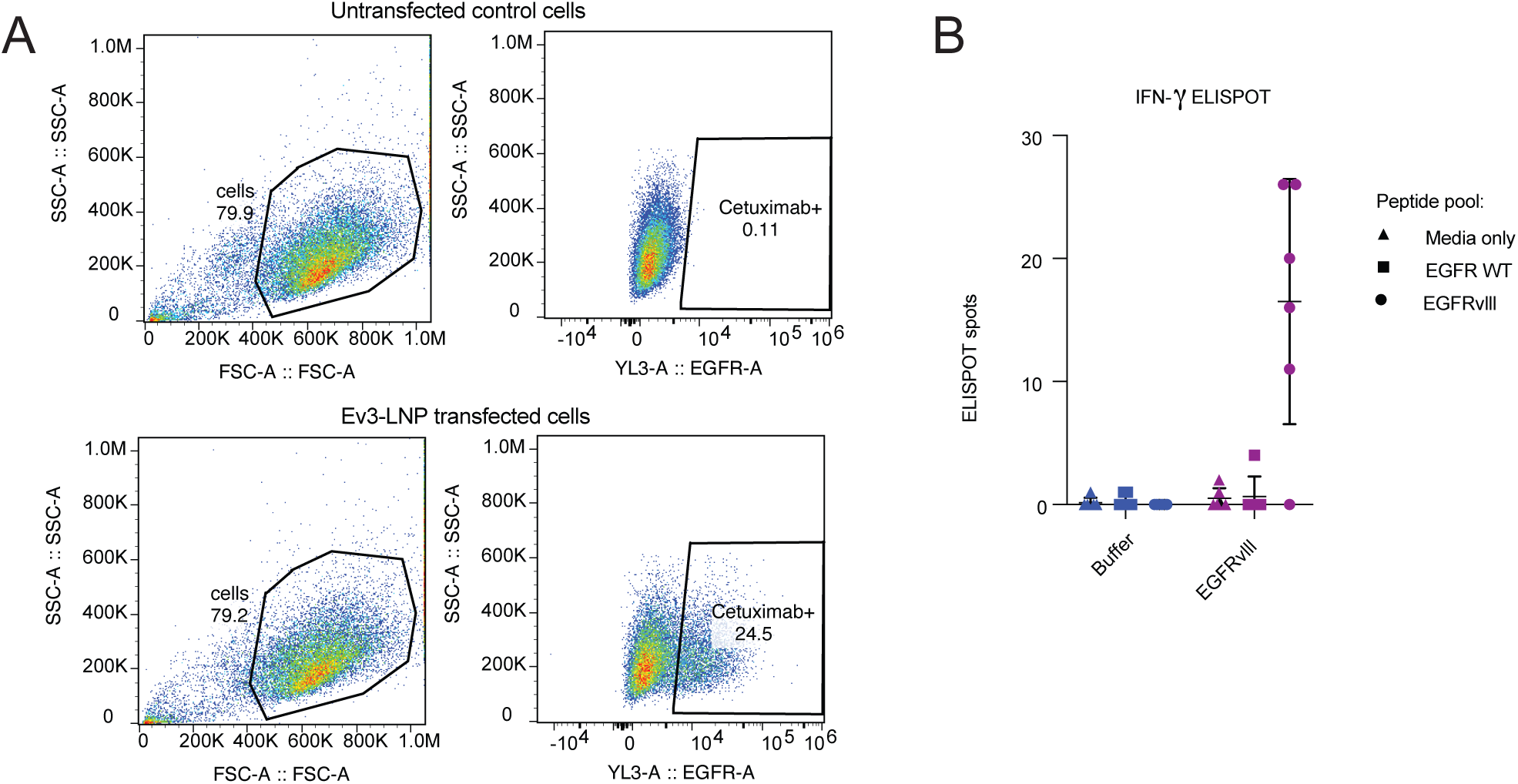
Surface EGFRvIII expression following mRNA-LNP transfection and antigen-specific IFN-γ responses. A) Flow cytometry gating strategy (left) and cetuximab staining (right) of untransfected control cells (top) and cells transfected with Ev3-LNP (bottom), confirming surface expression of EGFRvIII protein (0.11% vs 24.5% cetuximab+). B) IFN-γ ELISPOT of splenocytes collected at 21 dpi from buffer control- or Ev3-LNP-treated tumor-bearing mice, stimulated with media alone, a human EGFR wild-type peptide pool, or a peptide pool spanning the EGFRvIII exon 1-8 junction neoepitope.

**Supplementary Figure 3.**
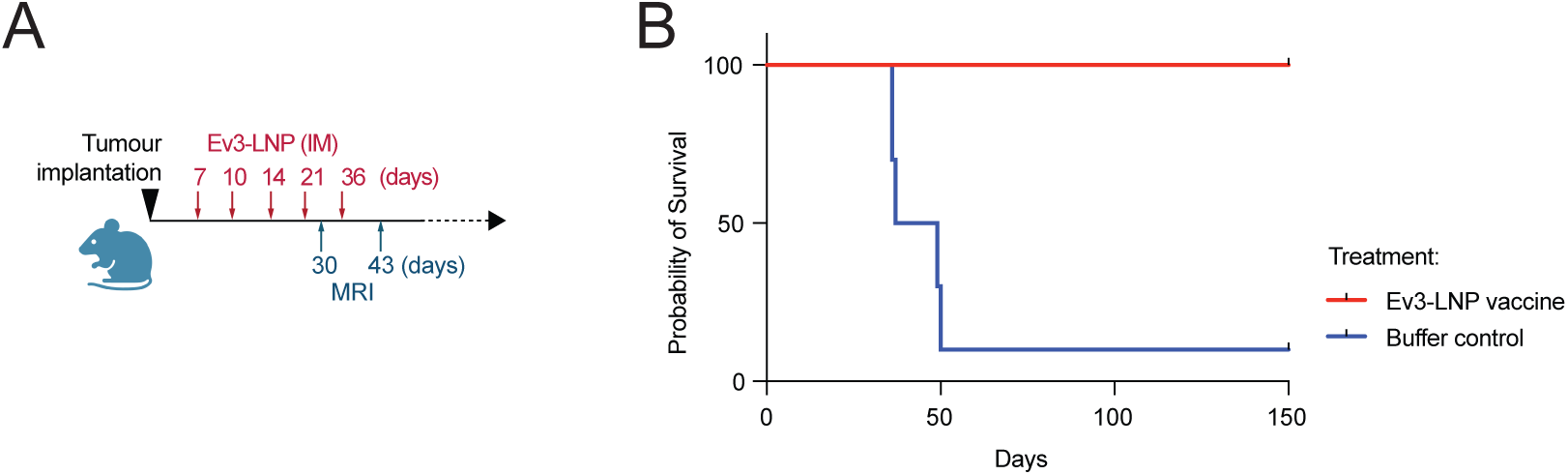
mRNA-LNP vaccination is efficacious in an independently generated male syngeneic model. a. Experimental design. Male C3H mice bearing orthotopic C3H-CL1-M1 tumors received Ev3-LNP intramuscularly at 7, 10, 14, 21 and 36 dpi, with MR imaging at 30 and 43 dpi. b. Corresponding Kaplan-Meier survival analysis of Ev3-LNP-versus buffer control-treated mice.

**Supplementary Figure 4.**
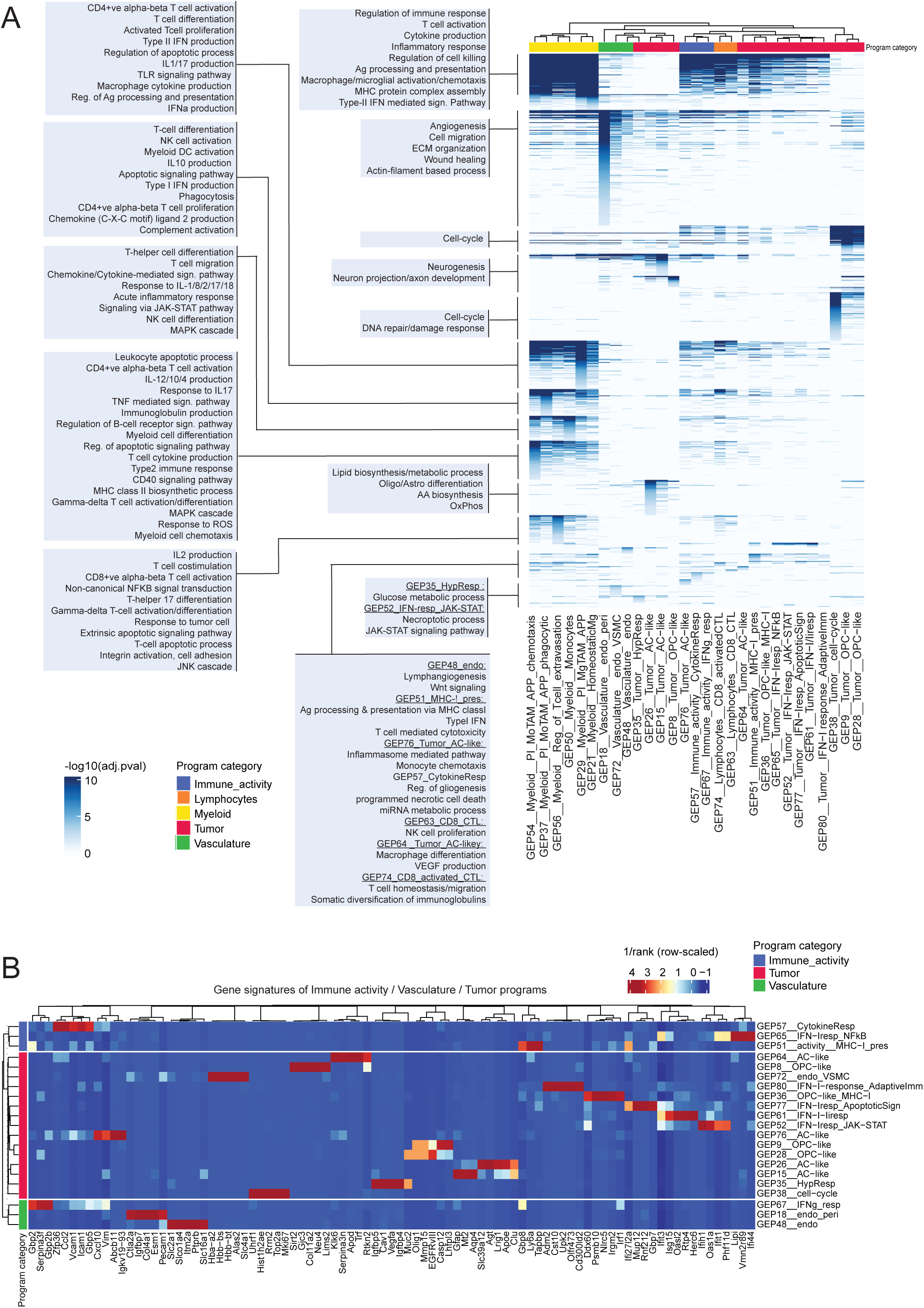
Gene signatures of lesion-enriched programs. A) Summary of biological processes (GO:BP; rows) significantly enriched (adjusted p-value < 0.05) among the top 2000 genes of lesion-enriched gene expression programs, ranked based on program gene scores. Enrichment of a BP is represented as -log10(adj.pvalue). B) Heatmap of inverse rank scores for the top five genes from immune activity, vasculature or tumor programs.

**Supplementary Figure 5.**
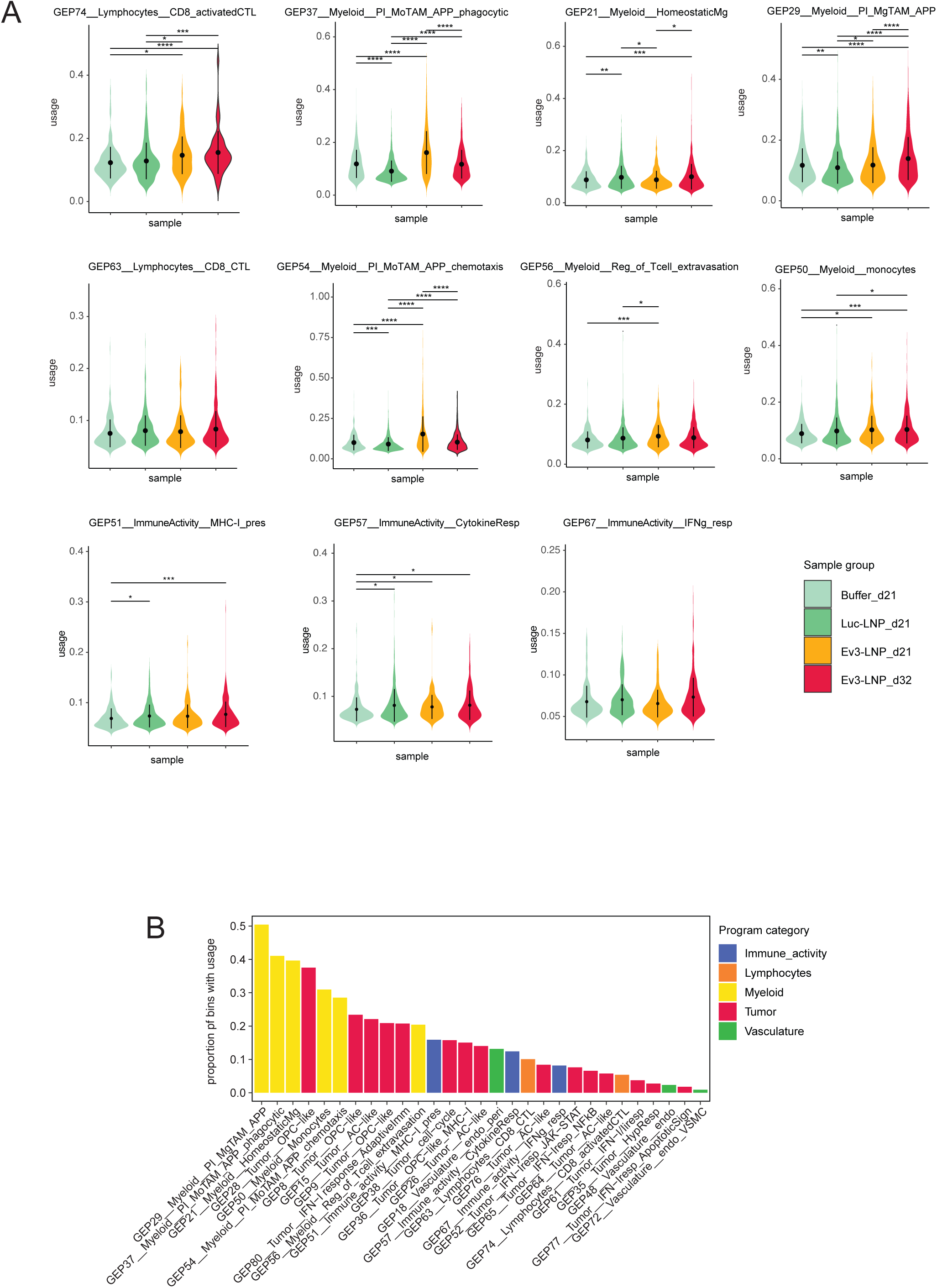
Usage distribution of lesion-enriched programs across the four experimental conditions. A) Violin plots showing the distribution of program usage for lymphocyte and selected myeloid programs within lesion bins across the four experimental conditions. Statistical significance was assessed using the Wilcoxon rank-sum test with Holm correction for multiple testing (ns: p > 0.05 (not reported in figure); *p ≤ 0.05; **p ≤ 0.01; ***p ≤ 0.001; ****p ≤ 0.0001). B) Barplot showing the proportion of bins with program usage > 0.05 across the cohort for each lesion-enriched program, ordered by decreasing prevalence and colored by the broad cell-type or activity class assigned to each program (Immune activity, Lymphocytes, Myeloid, Tumor or Vasculature).

**Supplementary Figure 6.**
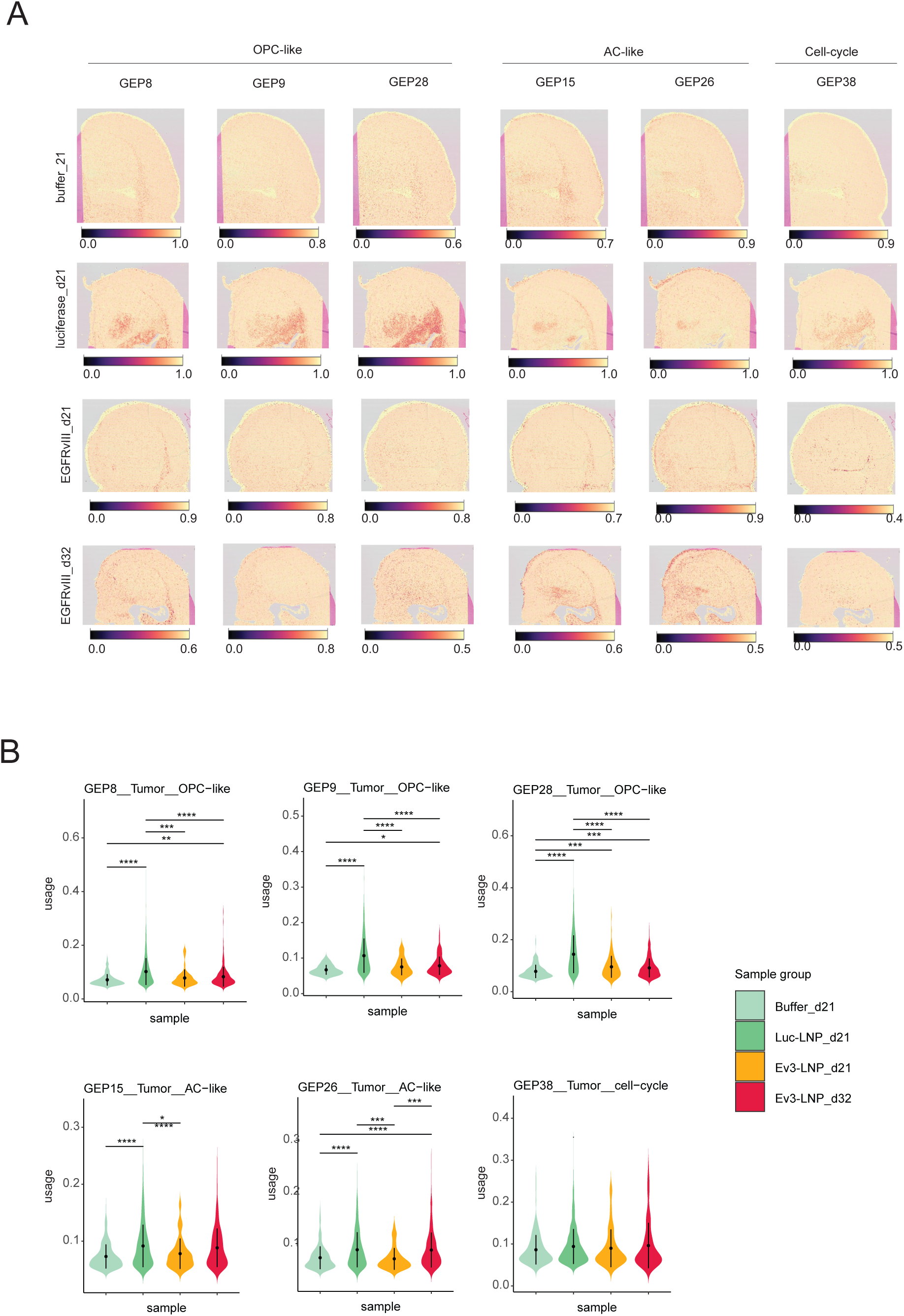
Gene expression programs of the tumor. A) Visium HD spatial plots showing usage of Tumor programs (OPC-like, AC-like and Cell-cycle) across tissue sections. B) Violin plots showing the distribution of program usage for tumor programs within lesion bins across the four experimental conditions. Statistical significance was assessed using the Wilcoxon rank-sum test with Holm correction for multiple testing (ns: p > 0.05 (not reported in figure); *p ≤ 0.05; **p ≤ 0.01; ***p ≤ 0.001; ****p ≤ 0.0001).

**Supplementary Figure 7.**
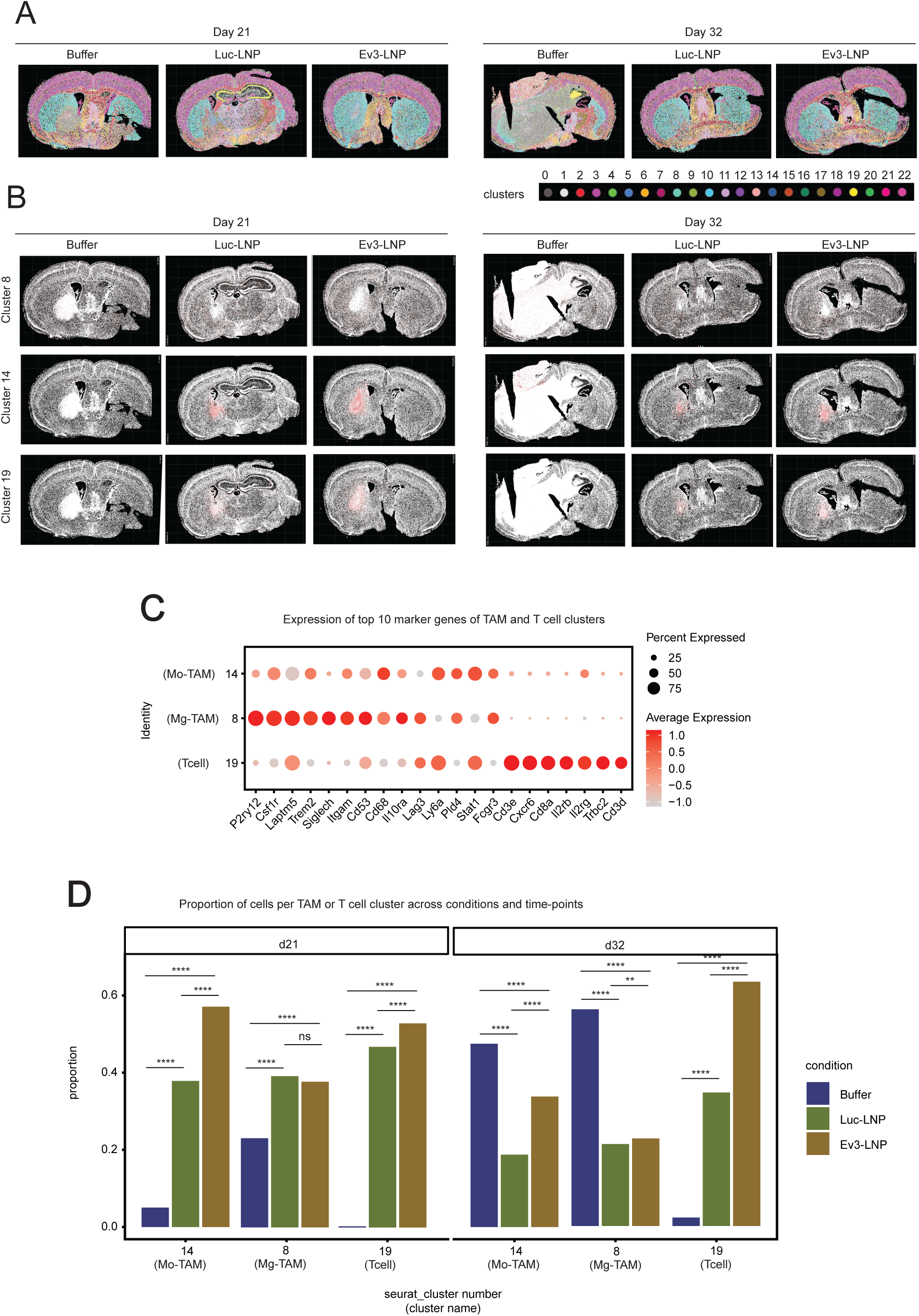
Xenium data analysis. A) Spatial plots of Seurat-derived Louvain clusters (row 1) in Xenium profiles of coronal mouse brain sections treated with buffer control, Luc-LNP or Ev3-LNP and sampled at 21 and 32 dpi. B) Spatial plots highlighting clusters 8 (Mg-TAM), 14 (Mo-TAM) and 19 (T cell) in each sample. C) Dotplot showing the expression of the top 10 marker genes per cluster, identified using Seurat’s FindAllMarkers() function. The dot size represents the percentage of cells within each cluster expressing the gene, and the dot color indicates scaled average expression. D) Grouped barplot showing the proportion of cells from each condition within clusters 8, 14 and 19, at each time-point. Statistical significance between conditions was assessed using a proportion test with Yates correction (ns: p > 0.05; *p ≤ 0.05; **p ≤ 0.01; ***p ≤ 0.001; ****p ≤ 0.0001).

**Supplementary Figure 8.**
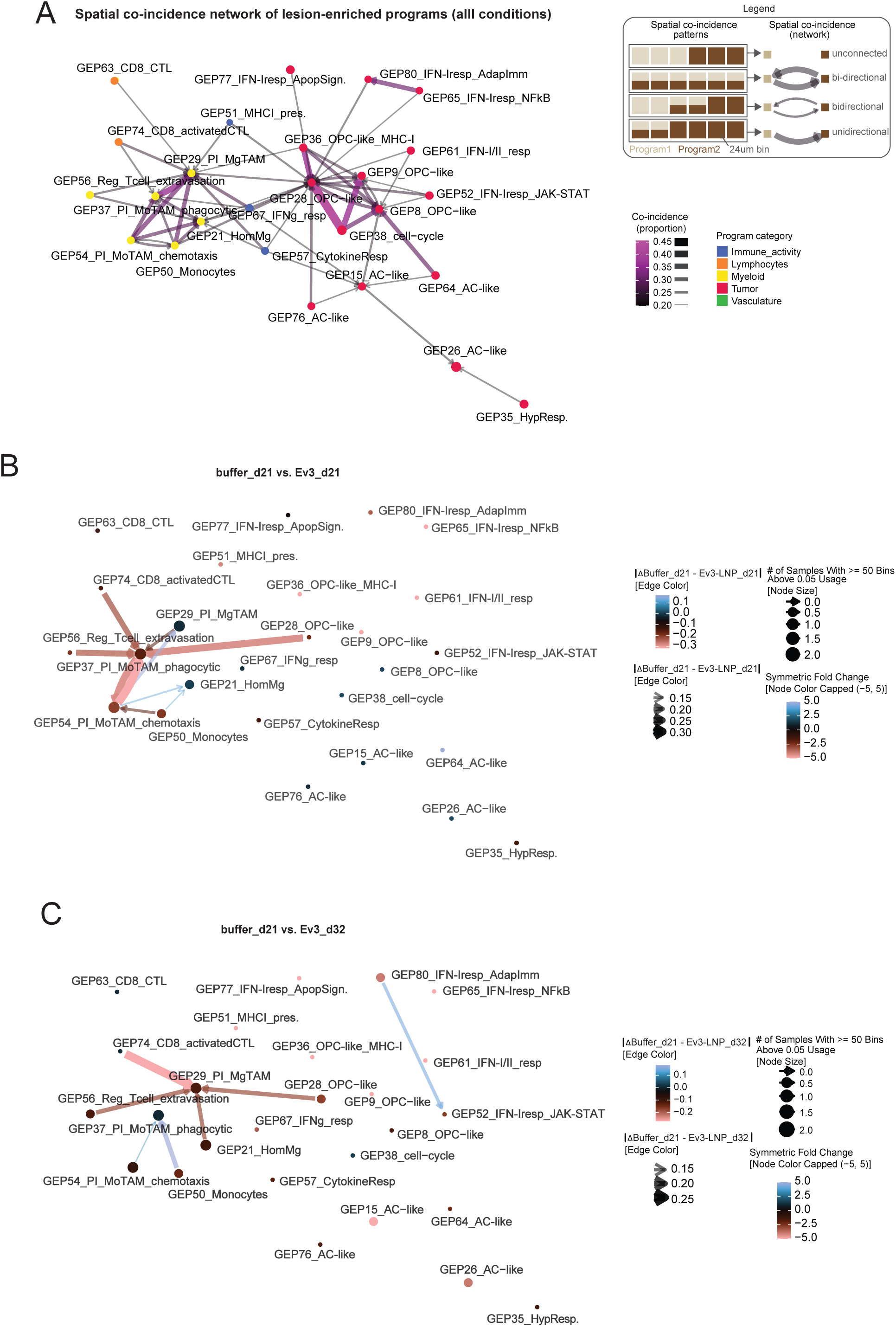
Spatial coincidence networks of lesion-enriched programs. A) Global spatial coincidence network of lesion-enriched programs across all conditions. Node color denotes the broad cell-type or activity class assigned to each program (Immune activity, Lymphocytes, Myeloid, Tumor or Vasculature); edge color and width represent the proportion of 24 μm bins in which the two programs co-occur, and edge style indicates whether co-incidence is uni-directional or bi-directional, as illustrated in the legend inset. B, C) Differential spatial coincidence networks comparing lesion-enriched programs in Buffer_d21 vs. Ev3-LNP_d21 (B) and Buffer_d21 vs. Ev3-LNP_d32 (C). Node size reflects the number of samples in which a query program has ≥50 bins above usage threshold >0.1, indicating its prevalence. Node color represents the symmetric fold change in proportion of program-positive bins between the two conditions, capped at ±5 to emphasize relative enrichment.

**Supplementary Figure 9.**
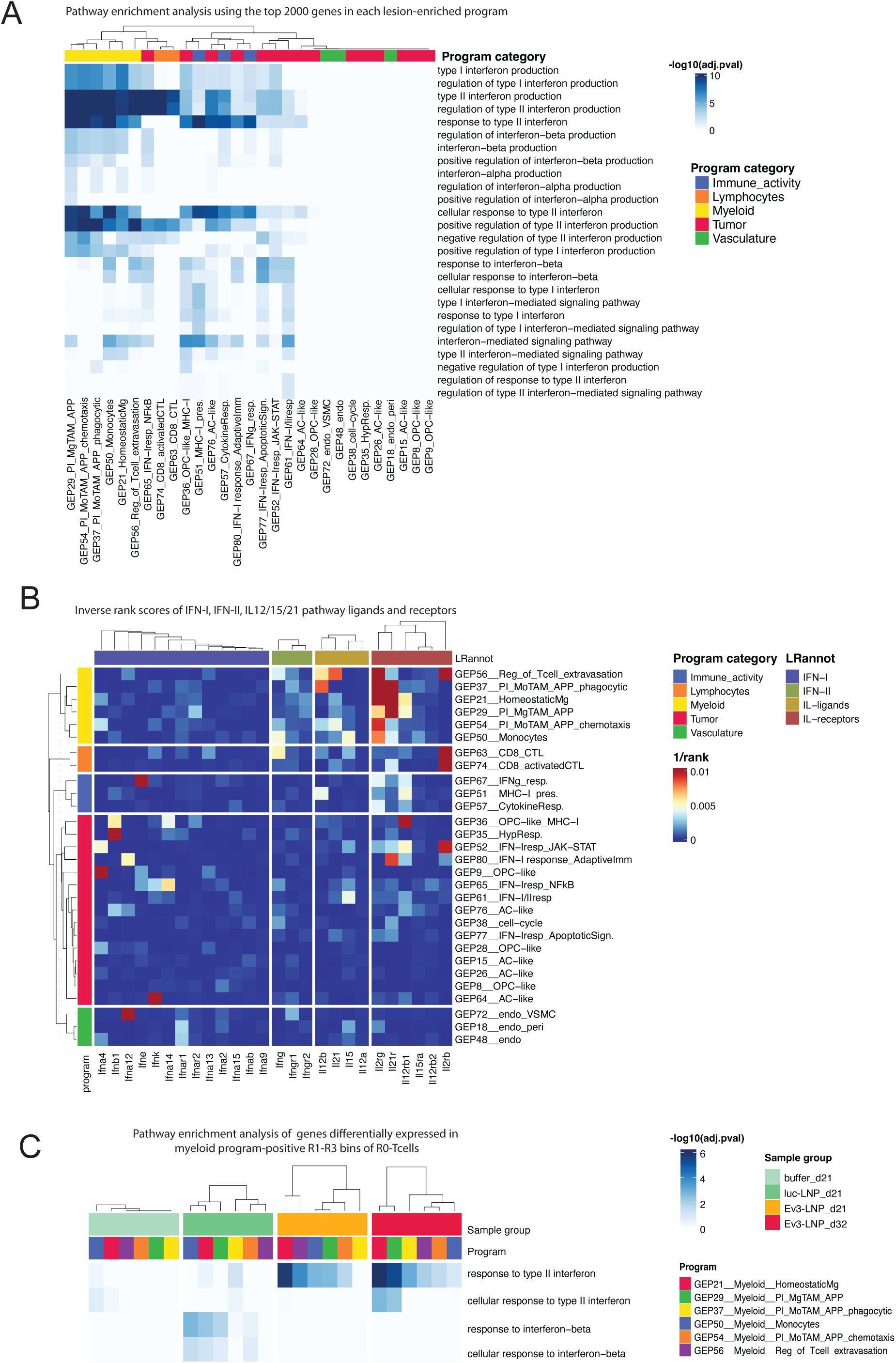
Interferon signaling. A) Summary of interferon signaling associated biological processes (GO:BP; rows) significantly enriched (adjusted p-value < 0.05) among the top 2000 genes of lesion-enriched gene expression programs, ranked based on program gene scores. Enrichment of a BP is represented as - log10(adj.pvalue). B) Heatmap of inverse rank scores of ligand and receptor genes from IFN-I, IFN-II, and IL12/15/21 pathways in lesion-enriched programs. Ligand and receptor genes were derived from CellChatDB. C) Pathway enrichment analysis using differentially expressed genes in myeloid program-positive bins in T cell neighbourhoods. Among the R1-R3 neighbours of the T cells, bins with > 0.05 usage of a given myeloid program were selected and compared to all other T cell neighbourhood bins (performed separately for each myeloid program and experimental condition). Significantly enriched genes per program-condition combination were subjected to GO Biological Process (GO:BP) enrichment analysis. The heatmap summarizes significantly enriched interferon signaling associated GO:BP terms (rows; adjusted p-value < 0.05), with enrichment represented as -log10(adjusted p-value).

**Supplementary Figure 10.**
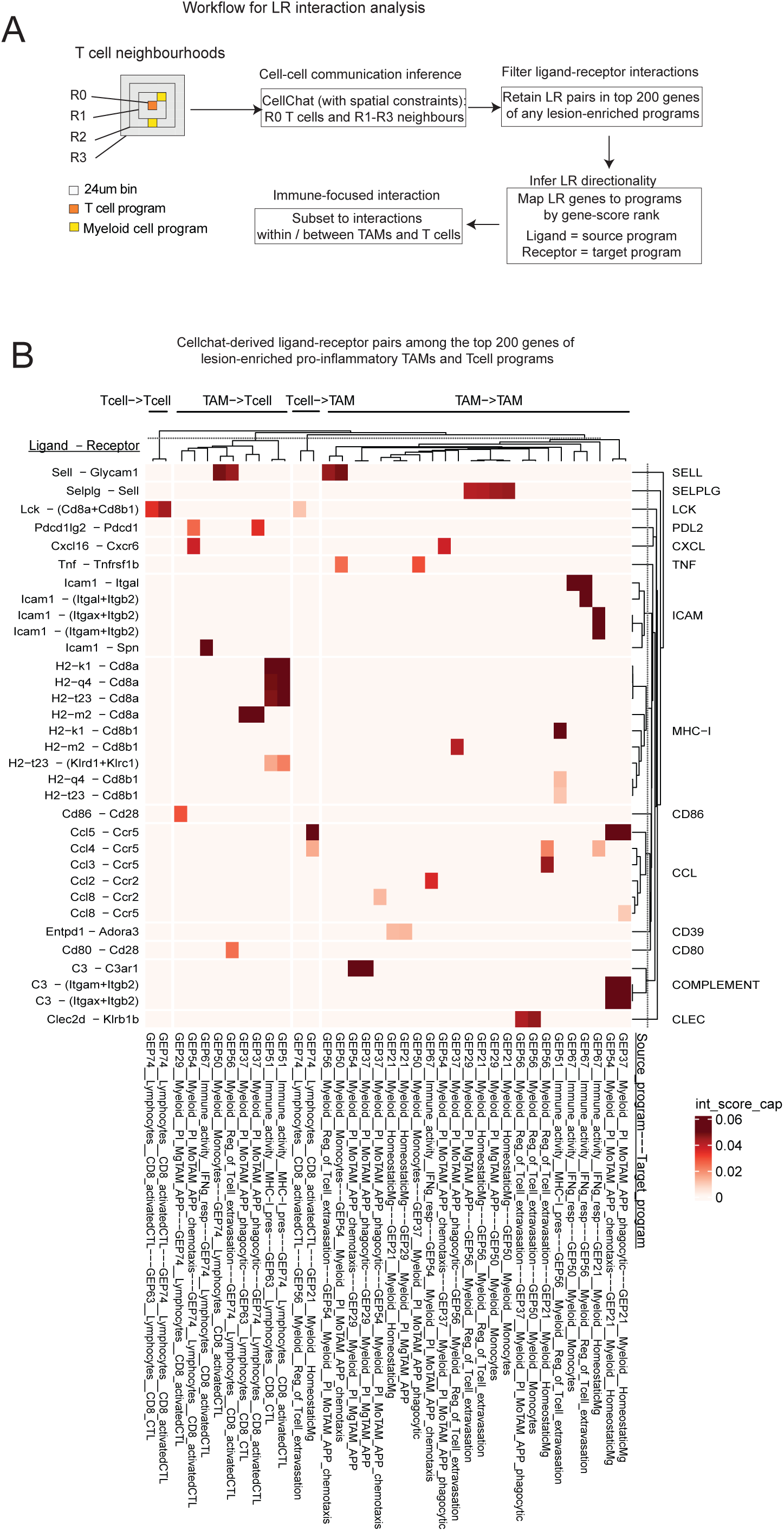
Determining T cell neighbourhoods and their ligand-receptor interactions. A) Flowchart describing the workflow for the ligand-receptor (LR) analysis: T cell neighbourhoods were defined around T cell program-positive index bins (R0) and their R1-R3 neighbours; cell-cell communication was inferred using CellChat with spatial constraints; LR pairs were retained only where both the ligand and the receptor ranked within the top 200 genes of any lesion-enriched program; directionality was inferred by assigning each gene to the program in which it achieved its highest gene-score rank (ligand = source program, receptor = target program); and interactions were subset to those occurring within or between TAM and T cell programs. B) CellChat-derived ligand-receptor pairs among the top 200 genes of lesion-enriched pro-inflammatory TAM and T cell programs. Heatmap of interaction scores of ligand-receptor pairs (y-axis) across pairs of myeloid and lymphocyte programs (source program — target program; x axis). Interaction scores were computed as the mean of gene ranks (derived from rank-ordered cNMF gene-scores) of the ligand in the source program and the receptor in the target program. LR interaction scores are reported only for program pairs in which both the ligand and receptor are present within the top 200 genes of their respective programs. The LR pairs are grouped by their corresponding pathway annotation from the CellChat mouse database. Program pairs are grouped based on their broad directional class (Tcell→Tcell, TAM→Tcell, Tcell→TAM and TAM→TAM).

## Table Legends

**Supplementary Table 1. Antibodies, stains and dextramers used in this study**

Reagents used for spectral flow cytometry, dextramer staining and immunostaining. The table lists, for each reagent, the target marker, fluorochrome or colour, clone, supplier, catalogue number and dilution factor, in three sections: (i) the 28 antibodies and viability stains used for spectral flow cytometry; (ii) the EGFRvIII H2-Kk PE dextramer (Immudex, custom order) used to identify anti-EGFRvIII CD8+ T cells by flow cytometry; and (iii) the primary antibodies used for immunohistochemistry and multiplexed immunofluorescence (EGFRvIII, Iba1, iNOS and Arg-1).

**Supplementary Table 2. Custom Visium HD probe sequences for EGFRvIII detection**

Sequences of the nine custom probe pairs designed to detect EGFRvIII transcripts in Visium HD spatial transcriptomic profiling. For each probe the table reports the probe identifier, the 25 bp left-hand side (LHS) and right-hand side (RHS) probe sequences targeting the EGFRvIII coding sequence, the partial Read 2S handle appended to the LHS probe, the 5′ phosphate and 30-mer poly(A) tail appended to the RHS probe, and the complete LHS and RHS sequences as ordered.

**Supplementary Table 3. Identification of lesion-enriched programs**

**a.** Wilcoxon test comparing program usage between lesion and non-lesion bins in each sample. For each program in a sample, bins with usage >0.05 were selected, and the distribution of usage values was compared between lesion and non-lesion bins using a Wilcoxon rank-sum test (see Fig. 3A and Supp. Fig. 5). The table reports, for each program: the mean usage in lesion and non-lesion bins in a sample, the log2 fold change of mean usage (log2[lesion / non-lesion]), and the associated p-value. **b.** Proportion test comparing program prevalence between lesion and non-lesion bins in each sample. Using prop.test() in R, the proportion of bins with usage >0.05 was compared between lesion and non-lesion regions in each sample. The table reports the total number of bins with usage >0.05 in a sample, the corresponding lesion and non-lesion bin counts, the proportions in each group, the log2 fold change of the proportions and the associated p-values.

